# More than a ligand: PD-L1 promotes oncolytic virus infection via a metabolic shift that inhibits the type I interferon pathway

**DOI:** 10.1101/2022.08.31.506095

**Authors:** Jonathan J. Hodgins, John Abou-Hamad, Ash Hagerman, Edward Yakubovich, Christiano Tanese de Souza, Marie Marotel, Ariel Buchler, Saleh Fadel, Maria M. Park, Claire Fong-McMaster, Mathieu F. Crupi, John C. Bell, Mary-Ellen Harper, Benjamin H. Rotstein, Rebecca C. Auer, Barbara C. Vanderhyden, Luc A. Sabourin, Marie-Claude Bourgeois-Daigneault, David P. Cook, Michele Ardolino

## Abstract

Targeting the PD-1/PD-L1 axis has transformed the field of immune-oncology. While conventional wisdom initially postulated that PD-L1 serves as the inert ligand for PD-1, an emerging body of literature suggests that PD-L1 has cell-intrinsic functions in immune and cancer cells. In line with these studies, here we show that engagement of PD-L1 via cellular ligands or agonistic antibodies, including those used in the clinic, potently inhibits the type I interferon pathway in cancer cells. Hampered type I interferon responses in PD-L1-expressing cancer cells resulted in enhanced infection with oncolytic viruses in vitro and in vivo. Consistently, PD-L1 expression marked tumor explants from cancer patients that were best infected by oncolytic viruses. Mechanistically, PD-L1 suppressed type I interferon by promoting a metabolic shift characterized by enhanced glucose uptake and glycolysis rate. Lactate generated from glycolysis was the key metabolite responsible for inhibiting type I interferon responses and enhancing oncolytic virus infection in PD-L1-expressing cells. In addition to adding mechanistic insight into PD-L1 intrinsic function and showing that PD-L1 has a broader impact on immunity and cancer biology besides acting as a ligand for PD-1, our results will also help guide the numerous efforts currently ongoing to combine PD-L1 antibodies with oncolytic virotherapy in clinical trials.

**Once sentence summary:** PD-L1 promotes oncolytic virus efficacy.

## INTRODUCTION

Being expressed on many cell types, PD-L1 is a readily available ligand for PD-1 present on immune cells in different tissues (*1*). The current model places PD-L1 as an agonistic ligand for PD-1, whose engagement results in inhibition of T and NK cells (*2*). This pathway is exploited by tumors as a mechanism of immune evasion, as evidenced by the success of antibodies blocking PD-1/PD-L1 interactions in several cancer indications (*3, 4*). However, the implementation of these therapies outpaced mechanistic understanding of this pathway, and many questions remain open on the PD-1/PD-L1 axis, including what other possible functions of PD-L1 are.

Emerging literature suggests that PD-L1, beyond its one-dimensional role as the ligand for PD-1, has cell-intrinsic signaling in cancer and immune cells (*5, 6*). PD-L1 signaling has been found to modulate many cellular processes, including the TGF-β pathway and the epithelial-mesenchymal transition (*7–10*), EGFR signaling (*11*), MAPK activation (*12, 13*), apoptosis (*14, 15*), DNA damage (*16–18*), proliferation and metastasis (*19*), and even cellular metabolism (*20, 21*). In most cases, the underlying mechanisms by which PD-L1 impacts these biological processes have not been uncovered.

One pathway shown to be regulated by PD-L1 is the type I interferon (IFN) pathway (*22–25*). Type I IFNs are a family of cytokines that induce a cellular anti-viral state via a JAK/STAT signaling pathway that promotes the transcription of hundreds of interferon-stimulated genes (ISGs) (*26*). Type I IFNs have other important roles in immunity and cell death (*27*), and it is in these contexts that their connection with PD-L1 has been established. However, the ability of PD-L1 to control viral infection, arguably the most prominent function of type I IFNs, has not been explored. When investigating PD-L1 expression in cancer cells, this relationship becomes even more important in light of the tremendous interest in combining PD-L1 blockade with Oncolytic Viruses (OVs).

OVs are viruses with a natural or engineered tropism for cancer cells over normal cells (*28, 29*), as a result of deficiencies in type I IFN signaling in cancer cells arising during transformation (*28, 29*). Preclinical and clinical studies frequently combine OVs with PD-1/PD-L1 blockade, or even engineer the OV to reduce PD-L1 expression in the tumor microenvironment (*30, 31*). However, the potential for synergy or antagonism between PD-L1 blockade and OVs should be carefully assessed prior to clinical translation.

To this end, we found that, by suppressing type I interferon responses, PD-L1 enhances infection with OVs in vitro and in vivo. Inhibition of type I IFNs depended on a metabolic shift promoted by PD-L1, resulting in enhanced rates of glucose uptake and glycolysis. Lactate generated from glycolysis was key to inhibit type I IFN responses. Taken together, our data mechanistically link PD-L1 cell intrinsic functions with susceptibility to OVs and provide a framework to further develop combinatorial therapies that better exploit the ability of PD-L1 to promote OV efficacy in cancer.

## RESULTS

### PD-L1 engagement promotes oncolysis of cancer cells

To test the hypothesis that PD-L1 regulates infection and oncolysis of cancer cells, we took advantage of the murine prostate cancer cell line TRAMP-C2 (*32*), which is widely used in OV preclinical studies (*33–35*), and constitutively expresses PD-L1 in culture (Figure S1A). To generate PD-L1-deficient cells, we targeted *Cd274* (the gene coding for murine PD-L1) by CRISPR/Cas9 and sorted cells lacking PD-L1 expression (Figure S1A).

To assess whether PD-L1 expression affects susceptibility to OVs, we infected parental and PD-L1-deficient TRAMP-C2 cells with the VSVΔ51, which is highly sensitive to the anti-viral effects of type I IFNs (*36, 37*). Strikingly, PD-L1 deletion resulted in a dramatic reduction of infection and OV-induced cell death (Figures 1A-C). To confirm that PD-L1 deletion, and not an experimental artefact, resulted in differences in OV infection, we complemented PD-L1 expression in TRAMP-C2-*Cd274*^-/-^ cells and tested if the phenotype was rescued. As expected, PD-L1 complementation enhanced VSVΔ51 infection compared to the empty vector control (Figures 1D-E). Increased resistance of PD-L1-deficient cells was observed not only in response to VSVΔ51, but also to vaccinia virus (Figure 1F), an oncolytic DNA virus undergoing clinical testing (*38*), showing that PD-L1 controls OV infection and oncolysis of cancer cells.

**Figure 1:**
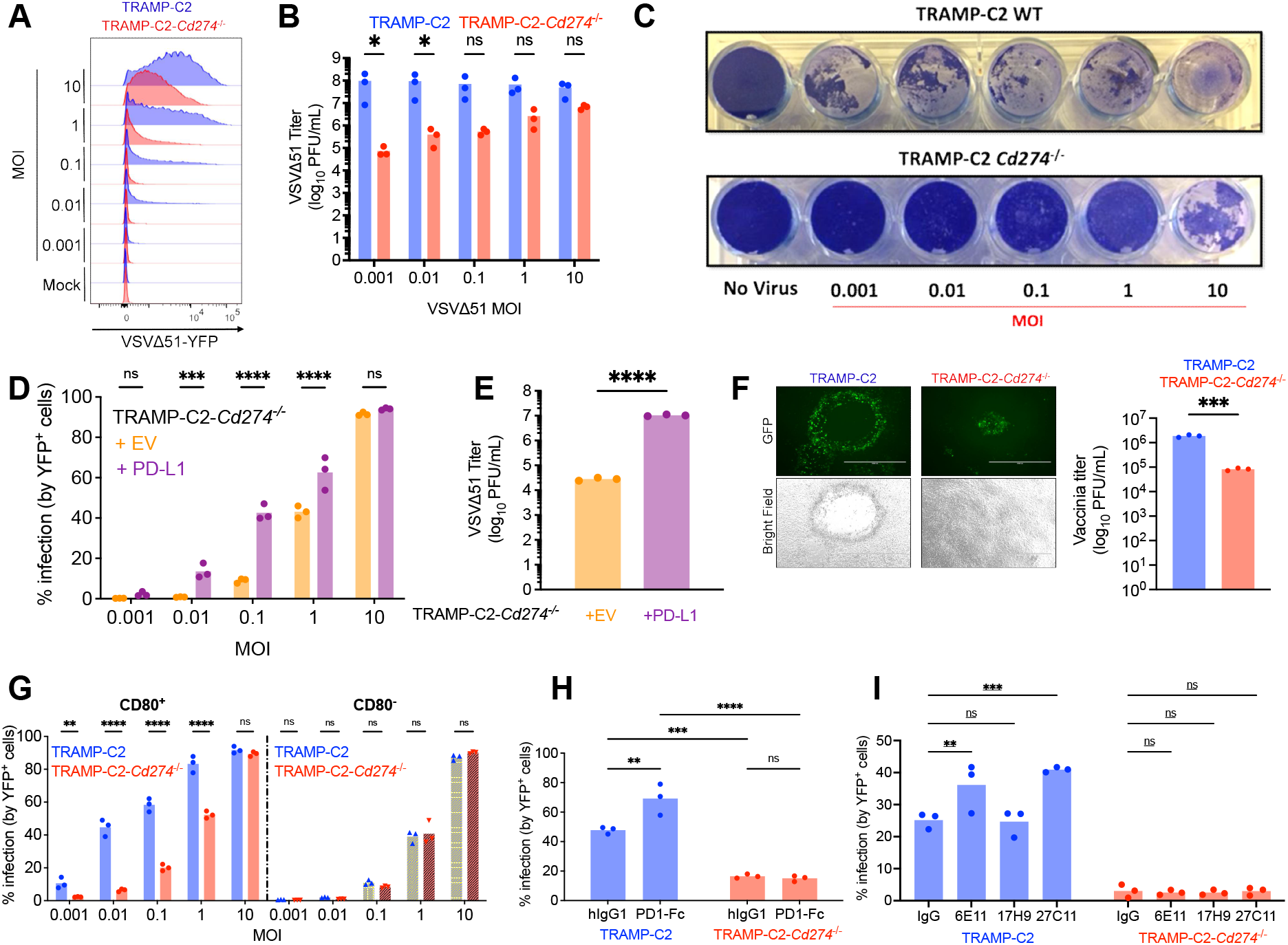
PD-L1 engagement promotes OV infection. (**A**-**C**) TRAMP-C2 and TRAMP-C2-*Cd274*^-/-^ cells were infected with VSV1′51-YFP at indicated MOIs and subjected to flow cytometry to quantify the viral YFP reporter 24 hours post-infection (A), plaque assay to quantify viral titers 24 hours post-infection (B), or Coomassie staining to visualize cell death 48 hours post-infection (C). The data depicted are representative of 4 experiments performed with similar results, n=3 biological replicates for the viral titer data. Statistical analysis by two-way ANOVA with Šídák’s correction for multiple comparisons. (**D-E**) TRAMP-C2-*Cd274*^-/-^ transduced with PD-L1 or empty vector were infected with VSV1′51-YFP at indicated MOIs for 24 hours and analyzed by flow cytometry **(D)**, or by plaque assay **(E)**. The data depicted are representative of 3 performed with similar results. Statistical analysis by two-way ANOVA with Šídák’s correction for multiple comparisons in **D** and two-tailed unpaired Student’s t-test in **E**. (**F**) TRAMP-C2 cells were infected with GFP-expressing vaccinia virus (Copenhagen strain) at MOI 0.1 for 48 hours, prior to fluorescence imaging. Representative of two performed with similar results. Viral titer was assessed by plaque assay, statistical analysis by two-tailed unpaired Student’s t-test (***: *p<0.001*). (**G**) CD80^+^ and CD80^-^ cells were isolated by FACS and subjected to VSV1′51-YFP at indicated MOIs for 24 hours prior to flow cytometry to quantify viral YFP reporter. The experiment depicted is representative of 3 performed with similar results. Statistical analysis by two-way ANOVA with Šídák’s correction for multiple comparisons. (**H**) TRAMP-C2 and TRAMP-C2-*Cd274*^-/-^ were pre-treated for 24 hours with 500 ng of recombinant PD-1-Fc chimeric protein, and infected with VSV1′51-YFP at MOI 0.1 for 24 hours prior to analysis by flow cytometry. The experiment depicted is representative of 3 performed with similar results. Statistical analysis by two-way ANOVA with Šídák’s correction for multiple comparisons. (**I**) TRAMP-C2 and TRAMP-C2-*Cd274*^-/-^ were pre-treated for 24 hours with 10 µg of PD-L1 antibodies, and infected with VSV1′51-YFP at MOI 0.1 for 24 hours prior to analysis by flow cytometry. The experiment depicted is representative of 3 performed with similar results. Statistical analysis by two-way ANOVA with Šídák’s correction for multiple comparisons.

Of the two known ligands for murine PD-L1 (*6*), TRAMP-C2 cells fail to express PD-1, whereas CD80 was expressed by ∼40% of cells in culture (Figure S1B). To determine if engagement by CD80 was needed for PD-L1 to enhance OV infection, we isolated CD80^+^ or CD80^-^ cells from both TRAMP-C2 or TRAMP-C2-*Cd274*^-/-^ by FACS and infected them with VSVΔ51. In the cells expressing CD80, PD-L1 enhanced viral infection (Figure 1G, left side). In stark contrast, in the absence of CD80, PD-L1 failed to promote viral infection and there was no longer any difference in infection between parental and PD-L1-deficient TRAMP-C2 cells (Figure 1G, right side). These data suggest that PD-L1 engagement is required to drive permissiveness to viral infection.

Given the abundance of PD-1 in the tumor microenvironment, we tested if PD-1 engagement of PD-L1 also resulted in an enhanced permissiveness to viral infection. Treatment with a recombinant PD-1-hIgG1 Fc chimeric protein enhanced infection in parental TRAMP-C2 cells compared to control-treated cells (Figure 1H), whereas PD-1-Fc treatment had no effect in PD-L1-deficient tumor cells (Figure 1H). These data show that both CD80 and PD-1 binding to PD-L1 promotes viral infection.

Next, we explored whether engagement of PD-L1 with antibodies promoted OV infection in cancer cells, a question of particular interest when considering the use of PD-L1 antibodies in the clinic. We pre-treated TRAMP-C2 with different PD-L1 antibodies characterized to prevent the interaction between PD-L1 and PD-1 (clone 27C11), PD-L1 and CD80 (clone 17H9), or both (clone 6E11) (*39*), which we reasoned may serve as agonists for PD-L1. Treatment with the two antibodies mimicking PD-1 binding to PD-L1, clones 27C11 and 6E11, significantly enhanced infection in parental TRAMP-C2, compared to isotype-treated, cells (Figure 1I), whereas no effect was observed with the clone mimicking CD80 interactions with PD-L1 (17H9), perhaps because CD80 was already present in our system. No effect was observed in PD-L1-deficient cells (Figure 1I). Taken together, this data confirms our hypothesis that PD-L1 engagement and signaling enhances susceptibility to infection. Moreover, the fact that direct engagement of PD-L1 with antibodies enhanced the susceptibility of cancer cells to infection rules out that uncharacterized signaling downstream of CD80 was responsible for the observed phenotype.

### PD-L1 promotes oncolysis by inhibiting type I IFN responses

Next, we set out to determine the mechanisms underlying PD-L1-driven enhancement of OV infection. We ruled out that entry of VSVΔ51 was impacted by deletion of PD-L1, as we found equal levels of viral RNA at early time points (e.g., 1-3 hours post-infection, Figure S2A), and both cell lines expressed similar levels of the VSV entry receptor LDL-R (Figure S2B).

As the type I IFN pathway is paramount for antiviral defence, we examined IFN responses induced by VSVΔ51 in parental and PD-L1-deficient cells. PD-L1-deficient TRAMP-C2 cells produced ∼2-fold more IFN-β compared to parental cells post-infection both at the protein and the transcript levels (Figures 2A-B) and, accordingly, had higher transcription of antiviral ISGs (Figure 2C), which was inhibited by PD-L1 re-complementation (Figure 2D and Figure S3A). PD-L1 regulated type I IFNs not only in response to viral infection, but also to the TLR3 and RIG-I agonist poly(I:C) (Figures S3B-C), suggesting that PD-L1 regulates the type I IFN response triggered by diverse stimuli, and corroborating other reports linking PD-L1 and regulation of type I IFNs (*22–25*).

**Figure 2:**
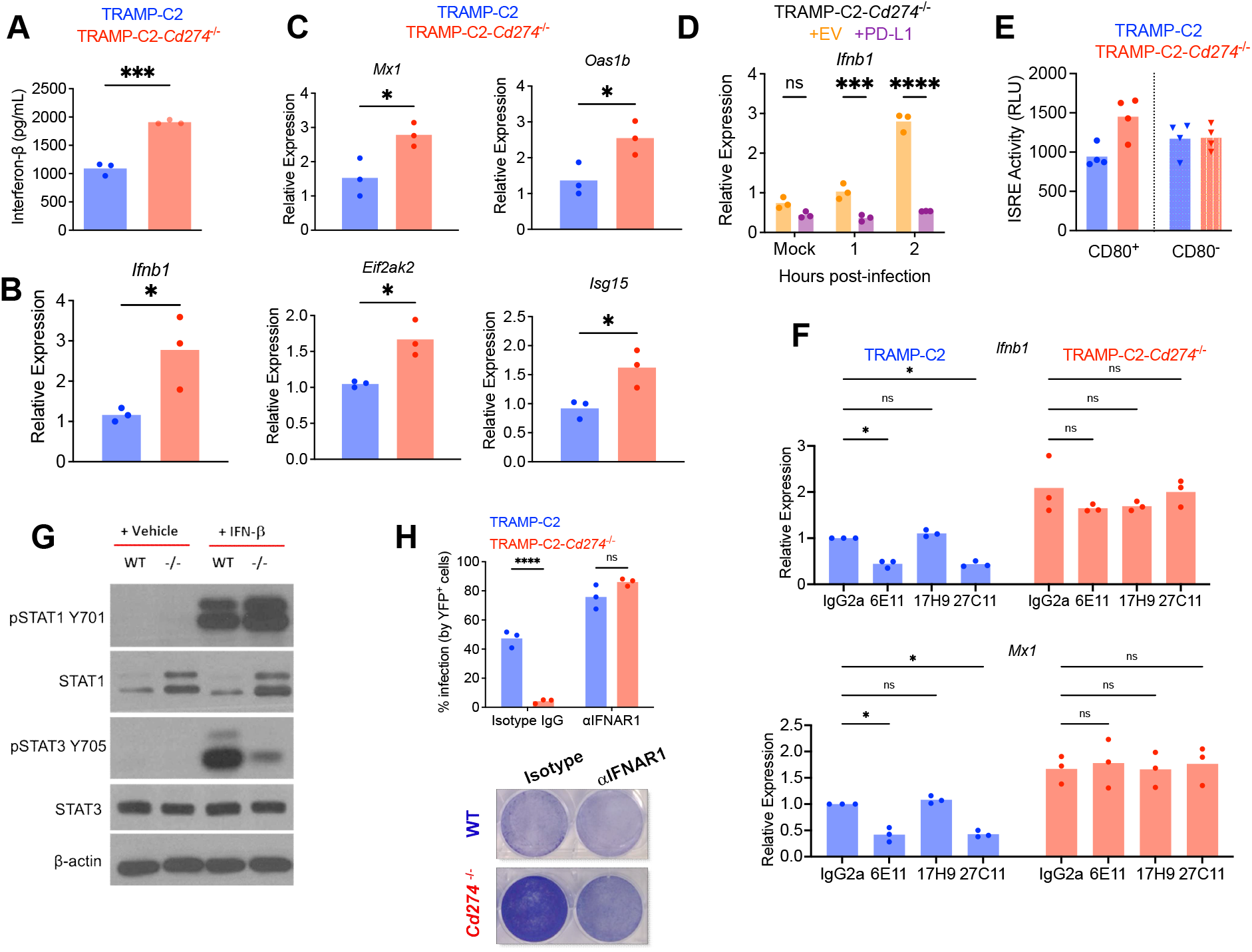
PD-L1 inhibits the type I IFN response to OVs. (**A-B**) IFN-β protein in culture supernatant (by ELISA, A) or transcript levels (by qPCR, B) were quantified following infection with VSV1′51-YFP for 8 hours. Statistical analysis with two-tailed unpaired Student’s t-test. (**C**) Expression of ISGs was quantified following infection with VSV1′51-YFP for 8 hours. Statistical analysis with two-tailed unpaired Student’s t-test. (**D**) TRAMP-C2-*Cd274*^-/-^ transduced with PD-L1 or empty vector were infected with VSV1′51-YFP at MOI 0.1 and analyzed by qPCR at indicated times post-infection to quantify *Ifnb1* transcripts. The data depicted are representative of 3 performed with similar results. Statistical analysis by two-way ANOVA with Šídák’s correction for multiple comparisons. (**E**) Type I IFN in culture supernatant was quantified using the L929-ISRE reporter cell line, in CD80^+^ and CD80^-^ cells isolated by FACS and infected with VSV1′51-YFP for 8 hours. The experiment depicted is representative of 3 performed with similar results. Statistical analysis by one-way ANOVA with Šídák’s correction for multiple comparisons. (**F**) TRAMP-C2 and TRAMP-C2-*Cd274*^-/-^ were pre-treated for 24 hours with 5 µg of PD-L1 antibodies, and infected with VSV1′51-YFP at MOI 0.1 for 8 hours prior to qPCR analysis. The experiment depicted is representative of 3 performed with similar results. Statistical analysis by two-way ANOVA with Šídák’s correction for multiple comparisons. (**G**) TRAMP-C2 and TRAMP-C2-*Cd274*^-/-^ cells were cultured in non-supplemented DMEM for 24 hours and treated with 200 units of recombinant murine IFN-β for 10 minutes prior to analysis by western blotting. The images depicted are representative of 3 performed with similar results. (**H**) TRAMP-C2 and TRAMP-C2-*Cd274*^-/-^ were pre-treated with 25 µg of anti-IFNAR1 for 24 hours, followed by infection with VSV1′51-YFP at MOI 0.1 for 24 hours prior to analysis by Coomassie staining or flow cytometry. The experiments depicted are representative of 3 performed with similar results. Statistical analysis by two-way ANOVA with Šídák’s correction for multiple comparisons.

PD-L1 engagement by CD80 was required to inhibit IFN-β production in TRAMP-C2 cells (Figure 2E). Additionally, antibodies cross-linking of PD-L1 further suppressed type I IFN responses in parental, but not PD-L1-deficient, TRAMP-C2 cells (Figure 2F). Taken together, these data show that PD-L1 engagement strongly dampens type I interferon responses.

We next assessed whether PD-L1, in addition to inhibiting IFN production, also controlled responsiveness to type I IFNs by regulating signaling downstream of their receptor. After acute stimulation of TRAMP-C2 cells with IFN-β we observed dramatic differences in signaling downstream of the type I IFN receptor; in particular, PD-L1 decreased the levels of STAT1 and promoted STAT3 phosphorylation at Tyr705 (Figure 2G). Re-expression of PD-L1 altered almost every step of the type I IFN pathway, from phosphorylation of TBK1 and IRF-3 (which together control initial production of type I IFN), to STAT1 levels and STAT3 phosphorylation (Figure S3D). Therefore, PD-L1 not only controls induction of type I IFNs, but also key signaling molecules involved in their responses.

To implicate type I IFNs as the pathway responsible for the PD-L1-mediated promotion of oncolysis, we pre-treated tumor cells for 24 hours with an antagonistic antibody against the type I IFN receptor subunit IFNAR1. Antibody treatment ablated the differences in infection between TRAMP-C2 and TRAMP-C2-*Cd274*^-/-^ cells (Figure 2H), suggesting that the phenotype was caused by PD-L1 inhibition of the type I IFN pathway. Altogether, these data show that PD-L1 engagement promotes oncolysis via inhibition of type I IFN responses.

### PD-L1 poises cancer cells to be more susceptible to OVs

The mechanistic link between PD-L1 promotion of OV infection and inhibition of type I IFN responses prompted us to hypothesize that parental and PD-L1-deficient TRAMP-C2 cells will be equally permissive to wild-type VSV infection, which, differently than VSVΔ51, blocks translation of newly synthesized type I IFNs upon infection. Surprisingly, PD-L1 still enhanced WT VSV infection of parental TRAMP-C2 cells compared to PD-L1-deficient TRAMP-C2 cells (Figure 3A), despite there being no detectable virus-induced type I IFN response in these cells (Figure S4).

**Figure 3:**
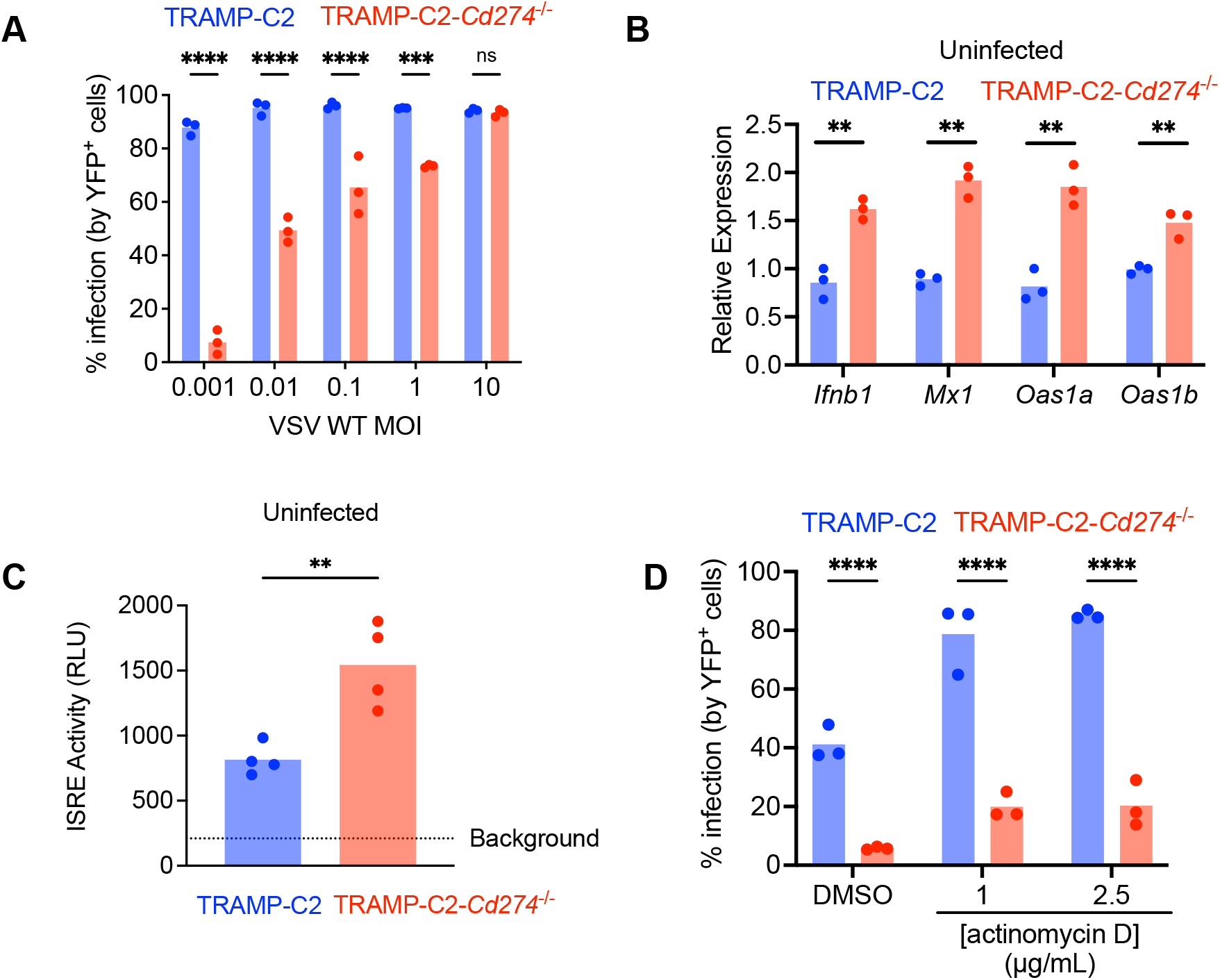
PD-L1 poises cancer cells to be more sensitive to viral infection. (**A**) TRAMP-C2 and TRAMP-C2-*Cd274*^-/-^ cells were infected with VSV WT at indicated MOIs for 24 hours prior to analysis by plaque assay. The experiment depicted is representative of 3 performed with similar results. Statistical analysis by two-way ANOVA with Šídák’s correction for multiple comparisons. (**B**) qPCR analysis of type I IFN and ISG transcripts expressed by TRAMP-C2 and TRAMP-C2-*Cd274*^-/-^ cells prior to infection. Statistical analysis by two-way ANOVA with Šídák’s correction for multiple comparisons. (**C**) Measurement of type I IFN in uninfected TRAMP-C2 and TRAMP-C2-*Cd274*^-/-^ supernatant using the L929-ISRE reporter line. The experiment depicted is representative of 3 performed with similar results. Statistical analysis by two-tailed unpaired Student’s t-test. (**D**) TRAMP-C2 and TRAMP-C2-*Cd274*^-/-^ cells were infected with VSV1′51-YFP at MOI 0.1 and treated with actinomycin D (or DMSO as vehicle control) at the time of infection. 24 hours later, cells were analyzed by flow cytometry. The experiment depicted is representative of 3 performed with similar results. Statistical analysis by two-way ANOVA with Šídák’s correction for multiple comparisons.

To better understand this unexpected result, we drew on literature showing that cancer cells often exhibit a constitutive type I IFN response (*40, 41*), and hypothesized that PD-L1 inhibited constitutive type I IFN responses. Indeed, TRAMP-C2 cells presented low but detectable expression of type I IFN and ISG transcripts even before infection, which was more pronounced in the absence of PD-L1 (Figure 3B). Using a sensitive IFN-reporter assay, we detected type I IFN activity in uninfected TRAMP-C2 culture supernatant, and more in PD-L1-deficient cells (Figure 3C). Therefore, TRAMP-C2 cells present a constitutive type I IFN response which is exacerbated by PD-L1 deletion.

The constitutive expression and regulation of type I IFNs, and the differential infection by VSV WT, suggested that PD-L1 poised cancer cells to a pro-viral state by constitutively repressing the type I IFN response and subsequent anti-viral ISG expression. To determine if this basal type I IFN response was sufficient to drive differences in OV infection, we treated TRAMP-C2 cells with actinomycin D at the time of VSVΔ51 infection to block all cellular transcription, including virus-induced production of type I IFNs. In this setting, the only source of type I IFNs comes prior to infection, as part of the constitutive IFN response observed in cancer cells. Despite the absence of virus-induced IFN, we still observed more VSVΔ51 infection in TRAMP-C2 compared to TRAMP-C2-*Cd274*^-/-^ cells (Figure 3D), suggesting that PD-L1 poises cancer cells to be more amenable to oncolysis.

### PD-L1 promotes a metabolic shift in cancer cells resembling the Warburg effect

To better understand the mechanisms underlying inhibition of type I IFNs by PD-L1, we performed RNA-sequencing on TRAMP-C2 and TRAMP-C2-*Cd274*^-/-^ cells, both before and after infection with VSVΔ51. We observed 2,690 and 5,486 differentially expressed genes in mock and infected samples respectively (FDR<0.05), confirming that PD-L1 regulated cellular pathways before infection (Figure S5A). To investigate the involvement of these pathways in the function of PD-L1, we experimentally activated or inhibited some of them and examined oncolytic viral infection in the presence and absence of PD-L1. TGF-β, EGFR and estrogen/androgen pathways were not found to be involved in the ability of PD-L1 to promote OV infection in cancer cells, as the phenotype was not lost upon experimental manipulation (Figures S5B-D).

Analysis of the differentially expressed genes showed that many metabolic enzymes were less abundantly expressed in PD-L1-deficient cells, resulting in a decreased glycolysis gene set score compared to parental TRAMP-C2 cells (Figure 4A), which was reflected in the differential Hypoxia pathway activity in the PROGENy analysis (Figure S5A). We observed increased expression of most glycolysis enzymes in TRAMP-C2 compared to TRAMP-C2-*Cd274*^-/-^ cells (Figure S6A), suggesting regulation of glycolysis by PD-L1. In corroboration to this hypothesis, we found that PD-L1-deficient cells had decreased glycolysis (Figure 4B) and increased oxidative phosphorylation (OXPHOS) (Figure 4C). Both phenotypes were fully rescued by re-expression of PD-L1 in TRAMP-C2-*Cd274*^-/-^ cells (Figures 4D-E). Accordingly, bioenergetic calculations (*42*) show that PD-L1 reduced ATP production from OXPHOS while increasing ATP production from glycolysis (Figures 4F-G). Many key parameters of mitochondrial respiration were reduced in PD-L1-expressing cells (Figure S6B) which also had reduced mitochondrial content (Figures S6C-D), without impacting mitochondrial ROS (Figure S6E). Increased glycolytic rate in parental TRAMP-C2 cells were also confirmed by untargeted metabolomics. We quantified approximately 100 water-soluble metabolites on TRAMP-C2 and TRAMP-C2-*Cd274*^-/-^ in the absence of infection (Table S1). The abundance of 52 metabolites was statistically different between parental and PD-L1-deficient cells (Fig. 4H) including the glycolysis metabolites fructose-1,6-biphosphate, dihydroxyacetone phosphate, and pyruvate, which are key indicators of glycolysis rates (*43*) (Figure S6F).

**Figure 4:**
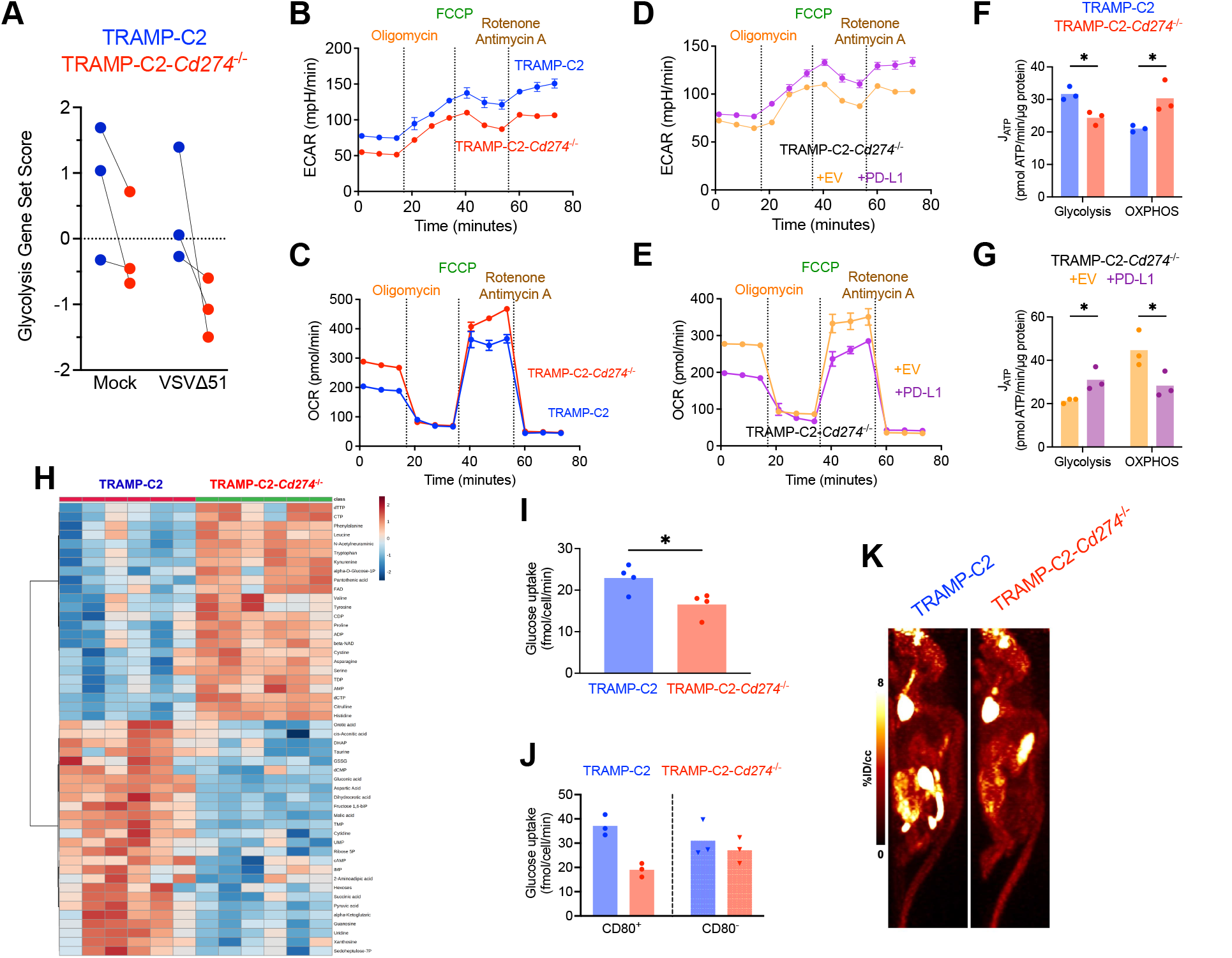
PD-L1 promotes Warburg metabolism in TRAMP-C2 cells. (**A**) Scoring for expression of genes included in the glycolysis gene set. RNA-seq dataset performed on TRAMP-C2 and TRAMP-C2-*Cd274*^-/-^ either mock-infected or infected with VSV1′51-YFP at MOI 0.1 for 8 hours (3 biological replicates per condition). **(B-E)** Seahorse metabolic flux analysis on TRAMP-C2, TRAMP-C2-*Cd274*^-/-^, TRAMP-C2-*Cd274*^-/-^ + EV, and TRAMP-C2-*Cd274*^-/-^ + PD-L1 (tested in n=6 technical replicates). Extracellular acidification rate (ECAR) and oxygen consumption rate (OCR) are measures of glycolysis and mitochondrial respiration, respectively. Images depicted are representative of 3 experiments with similar results. (**F-G**) Bioenergetic calculations based on Seahorse metabolic flux assay. N=3 biological replicates. **(H)** Hierarchical clustered heat map depicting relative abundance of 50 metabolites quantified in untargeted metabolomics of TRAMP-C2 and TRAMP-C2-*Cd274*^-/-^ cells. 6 biological replicates per cell line are shown. **(I-J)** Glucose uptake was measured in TRAMP-C2 and TRAMP-C2-*Cd274*^-/-^ cells (I), as well as CD80^+^ vs CD80^-^ cells FACS-isolated from TRAMP-C2 or TRAMP-C2-*Cd274*^-/-^ cells (J). The experiments depicted are representative of 3 performed with similar results. Statistical analysis by two-way ANOVA with Šídák’s correction for multiple comparisons. **(K)** Male NCG mice were implanted with subcutaneous TRAMP-C2 or TRAMP-C2-*Cd274*^-/-^ tumors. Standardized uptake value (SUV) of [^18^F]-fluorodeoxyglucose was assessed by PET imaging. The two mice shown are representative of 5-6 analyzed.

Not only were PD-L1 expressing cells more glycolytically active, but they also presented enhanced rate of in vitro glucose uptake (Figure 4I), which was dependent on PD-L1 engagement by CD80 (Figure 4J). The same phenotype was conserved in vivo, as determined in subcutaneous TRAMP-C2 and TRAMP-C2-*Cd274*^-/-^ tumors established in immunodeficient NCG mice subjected to PET scanning with the radiolabeled glucose analog [^18^F]-fluorodeoxyglucose. In this model, any differences would be driven by PD-L1 activity on cancer cells, rather than immune-dependent or PD-1-dependent mechanisms. Parental and PD-L1-deficient tumors grew at similar rates in NCG mice. Consistent with our in vitro data, parental TRAMP-C2 tumors had enhanced rates of [^18^F]-fluorodeoxyglucose uptake compared to PD-L1-deficent tumors (Figures 4K and Fig. S6G). Higher rates of glycolysis and glucose uptake, along with increased reliance on glycolysis for ATP generation, are highly consistent with the Warburg effect, where cancer cells preferentially use glycolysis over OXPHOS to meet bioenergetic demands and generate other metabolites (*44*).

### PD-L1 promotes glycolysis and inhibits IFN responses in human cancer cells

If PD-L1 regulation of type I IFN is a well-conserved feature, we expect our findings to be replicated in other cancer cell lines with similar features, in particular the metabolic characteristics of TRAMP-C2 cells. To test this hypothesis, we made use of an RNA-seq dataset of 675 human cancer cell lines (*45*) and scored each of those cell lines for their expression of PD-L1 (*CD274*) and expression of genes in the Glycolysis gene set, which includes glycolysis and other metabolic enzymes (Figure 5A). We hypothesized that in cell lines with high PD-L1 expression and high score for the Glycolysis gene set, PD-L1 would promote glycolysis, inhibit type I IFN responses and make tumor cells more susceptible to OVs. From this analysis, we chose two readily available cell lines: the renal cell carcinoma line 786-0 and the gastric carcinoma line Hs746, both with high glycolysis gene set scores, but with different levels of PD-L1. In both cell lines, we deleted PD-L1 using CRISPR/Cas9 (Figure S7A), and subjected parental and *CD274*^-/-^ cells to VSVΔ51 infection. In both 786-0 and Hs746 cells, PD-L1 deletion made tumor cells more resistant to OV (Figure 5B-C), consistent with our hypothesis and the data in the TRAMP-C2 model. Furthermore, PD-L1 inhibited the constitutive and virus-induced interferon response, as well as signaling downstream of the type I IFN receptor in both cell lines (Figure 5D-E and Figures S7B-C).

**Figure 5:**
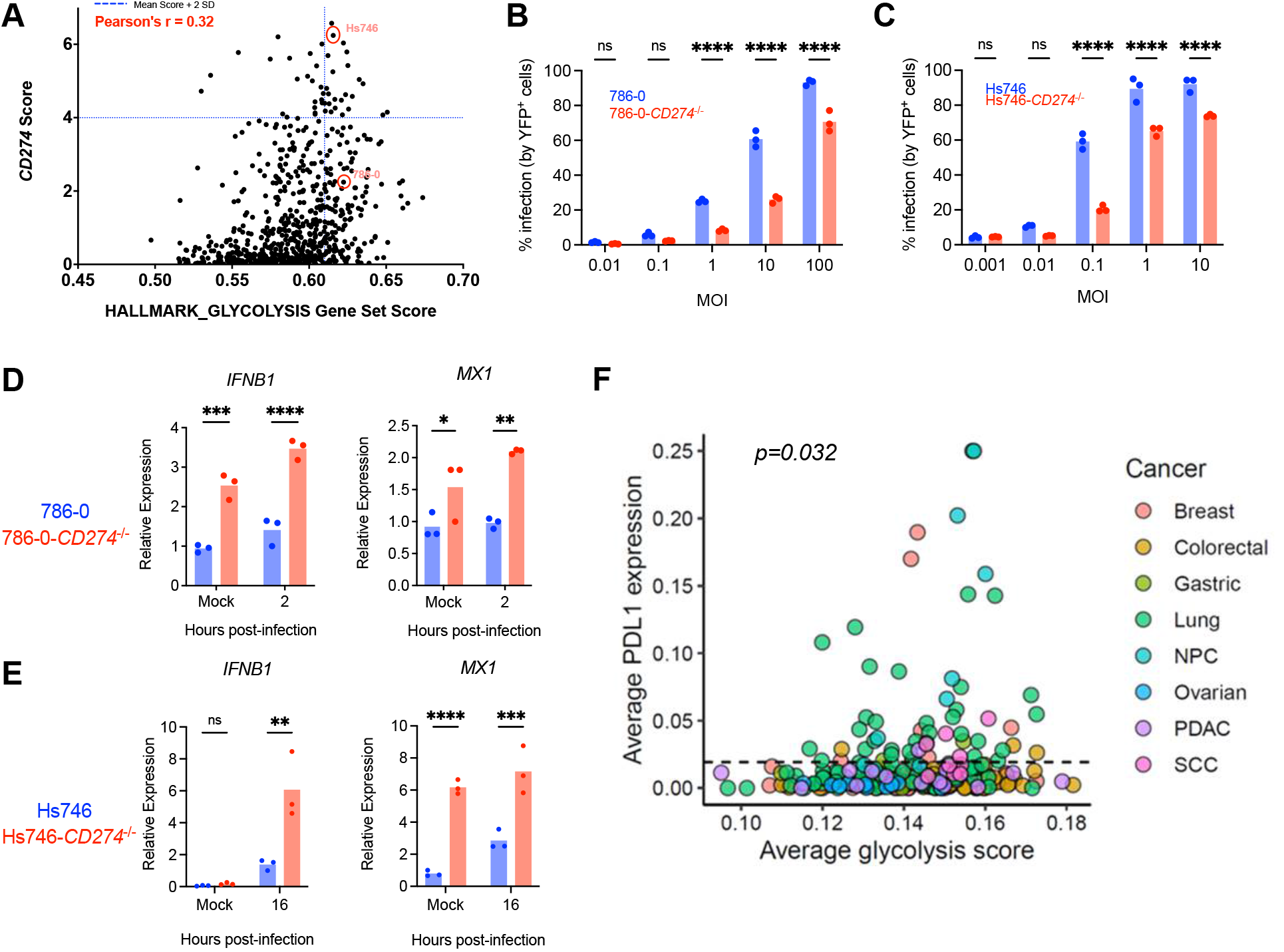
PD-L1 inhibits type I IFN responses in human cancer cells. (**A**) Bioinformatic analysis of publicly available RNA-seq from 675 human cancer cell lines, scored for their expression of PD-L1 (*CD274*) and glycolysis gene signatures. Blue lines on plot indicate mean + 2 SD; Pearson correlation coefficient is indicated. (**B** and **C**) 786-0 and Hs746 (WT and *CD274*^-/-^) infected with VSV1′51-YFP at indicated MOIs for 24 hours prior to analysis by flow cytometry. Experiments depicted are representative of 3 performed with similar results. Statistical analysis by two-way ANOVA with Šídák’s correction for multiple comparisons. (**D-E**) 786-0 and 786-0-*CD274*^-/-^ cells (D) or Hs746 and Hs746-*CD274*^-/-^ cells (E) infected with VSV1′51-YFP at MOI 1 or 0.1, respectively (or mock-infected) prior to qPCR analysis for *IFNB1* and *MX1* transcripts. n=3 biological replicates. Statistical analysis by two-way ANOVA with Šídák’s correction for multiple comparisons. *: p<0.05; **: p<0.01; ***: p<0.001; ****: p<0.0001. (**F**) Association between the expression of PD-L1 and *Glycolysis*-associated genes in the malignant cells of 266 tumours. Each point represents the average profile of malignant cells from scRNA-seq data sets, and dotted line represents mean PD-L1 expression of samples. Expression values reflect log-transformed gene counts per 10k transcripts and *Glycolysis* activity represents gene set scores from the associated MSigDB Hallmark gene set. Statistical significance assessed by Wilcoxon rank-sum test to compare glycolysis scores of tumors expressing PD-L1 above or below mean expression. NPC: nasopharyngeal cancer, PDAC: pancreatic ductal adenocarcinoma, SCC: squamous cell carcinoma.

To test if the link between PD-L1 expression and glycolysis in cancer cells held true in primary human samples, we took advantage of a single-cell RNA-sequencing dataset of 266 human tumors, spanning 8 types of cancer (Table S2). We scored each tumor for expression of PD-L1 and genes included in the glycolysis gene set. When we correlated the glycolysis score to PD-L1 expression at the single cell level in these cancers (*46*), we observed that tumors with high expression of PD-L1 had significantly higher glycolysis gene scores, in line with our hypothesis that PD-L1 drives glycolytic metabolism in cancer cells (Figure 5F). Therefore, the effect of PD-L1 on OV infection, type I IFN responses, and cellular metabolism is not unique to the TRAMP-C2 model and is conserved in other human cancer cell types.

### PD-L1 inhibits type I IFN via lactate dynamics

Metabolic alterations are now known to control inflammatory pathways, including type I IFN (*47*), e.g., glycolytic enzymes and metabolites are key regulators of inflammatory cytokines and anti-viral defenses (*48–51*). In line with this literature and considering our data, we reasoned that PD-L1 inhibition of type I IFN responses was linked with its ability to promote glycolysis. Recent research has mechanistically linked lactate to regulation of the type I IFN response (*52*). Lactate is an important metabolite generated during Warburg metabolism, whose physiological role is now beginning to be uncovered (*53*). Given its emerging role as a regulator of inflammatory responses, we hypothesized that lactate was responsible for the ability of PD-L1 to inhibit type I IFNs and promote virus infection.

In corroboration to our hypothesis, lactate was more abundantly produced in parental, over PD-L1-deficient cancer cells lines (Figure 6A). To directly test the role of lactate in PD-L1-driven inhibition of type I IFN responses, we pharmacologically perturbed lactate abundance prior to VSVΔ51 infection. First, to suppress lactate production, we used sodium oxamate and GNE-140: two structurally distinct inhibitors of the enzymes responsible for conversion of pyruvate into lactate: lactate dehydrogenases. Both inhibitors blocked the ability of PD-L1 to enhance virus infection (Figures 6B-C). On the other hand, treatment with lactate increased the permissiveness of PD-L1-deficient cells to OV infection, phenocopying the effect of PD-L1 (Figure 6D). Further, boosting glycolysis and lactate production through treatment with the ATP synthase inhibitor oligomycin also mimicked the effect of PD-L1 on VSVΔ51 infection (Figure 6E). We observed changes in susceptibility to OVs when tampering with lactate abundance not only in TRAMP-C2 cells (Figure 6A-E), but also in 786-0 and Hs746 cells (Figures 6F-I). Taken together, these experiments highlight the key role of lactate in promoting PD-L1-driven permissiveness to OV infection.

**Figure 6:**
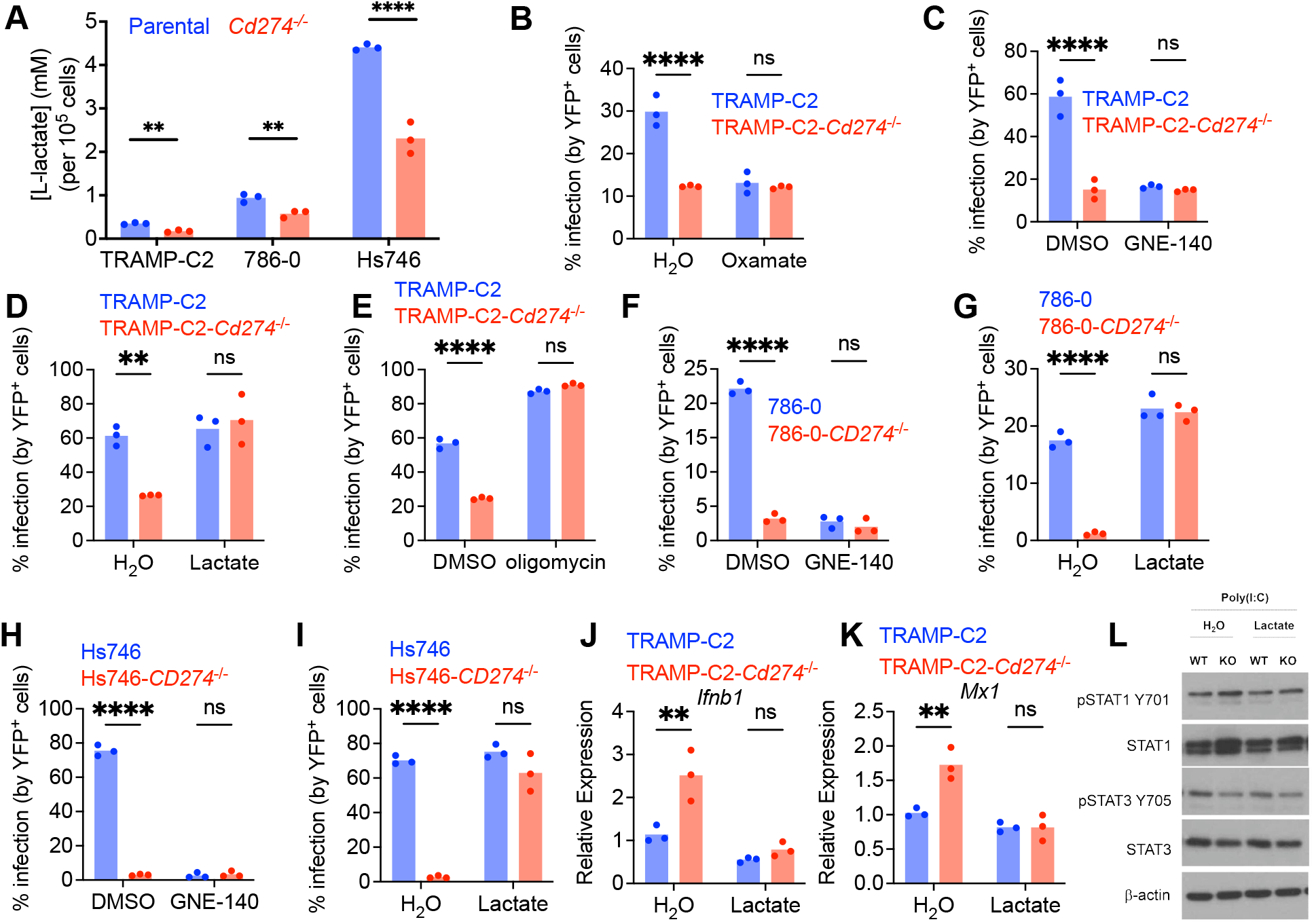
PD-L1 inhibits type I IFN via lactate dynamics. (**A**) Lactate quantification in TRAMP-C2, 786-0 or Hs746 culture supernatant. n=3 biological replicates. Statistical analysis by two-tailed unpaired Student’s t-test. *:p<0.05; ***: p<0.001 (**B-E**) TRAMP-C2 and TRAMP-C2-*Cd274*^-/-^ cells were pre-treated with oxamate at 10 mM (B); GNE-140 at 10 µM (C); lactate at 5 mM (D); oligomycin at 0.2 µM (E) for 24 hours, followed by infection with VSV1′51-YFP at MOI 0.1 for 24 hours prior to analysis by flow cytometry. Experiments depicted are representative of 3 performed with similar results. Statistical analysis by two-way ANOVA with Šídák’s correction for multiple comparisons. (**F-G**) 786-0 and 786-0-*CD274*^-/-^ cells were pre-treated with GNE-140 at 10 µM (F) or lactate at 5 mM (G) for 24 hours, followed by infection with VSV1′51-YFP at MOI 1 for 24 hours prior to analysis by flow cytometry. Experiments depicted are representative of 3 performed with similar results. Statistical analysis by two-way ANOVA with Šídák’s correction for multiple comparisons. (**H-I**) Hs746 and Hs746-*CD274*^-/-^ cells were pre-treated with GNE-140 at 10 µM (H) or lactate at 5 mM (I) for 24 hours, followed by infection with VSV1′51-YFP at MOI 0.1 for 24 hours prior to analysis by flow cytometry. Experiments depicted are representative of 3 performed with similar results. Statistical analysis by two-way ANOVA with Šídák’s correction for multiple comparisons. (**J-K**) TRAMP-C2 and TRAMP-C2-*Cd274*^-/-^ cells were pre-treated with lactate at 5 mM for 24 hours, followed by infection with VSV1′51-YFP at MOI 0.1 for 8 hours prior to qPCR analysis for *Ifnb1* and *Mx1*. n=3 biological replicates. Statistical analysis by two-way ANOVA with Šídák’s correction for multiple comparisons. (**L**) TRAMP-C2 and TRAMP-C2-*Cd274*^-/-^ cells were pre-treated with lactate at 5 mM for 24 hours, followed by transfection with poly(I:C) for 6 hours prior to western blotting analysis. Images depicted are representative of 3 with similar results.

To associate the PD-L1-mediated metabolic switch with the type I IFN response, we examined IFN-β induction and IFN receptor signaling following treatment with lactate. Lactate treatment resulted in normalization of the type I IFN response between parental and PD-L1-deficient TRAMP-C2 cells (Figures 6J-L), indicating that lactate is responsible for PD-L1-mediated regulation of type I IFNs.

### PD-L1 promotes OV infection in vivo

To determine if PD-L1 retains its ability to enhance OV infection in the more complex tumor microenvironment, we investigated whether PD-L1 promoted cancer cell infection in vivo. We established subcutaneous TRAMP-C2 or TRAMP-C2-*Cd274*^-/-^ tumors in immunodeficient NCG mice and, when tumors reached ∼750 mm^3^, we injected them with VSVΔ51 (expressing a luciferase reporter). After 24 hours we assessed viral infection by in vivo imaging and plaque assays. In corroboration of our in vitro studies, PD-L1-deficient tumors presented reduced infection compared to parental tumors (Figures 7A-B), indicating that PD-L1 expression on tumor cells drives increased OV infection in vivo.

**Figure 7:**
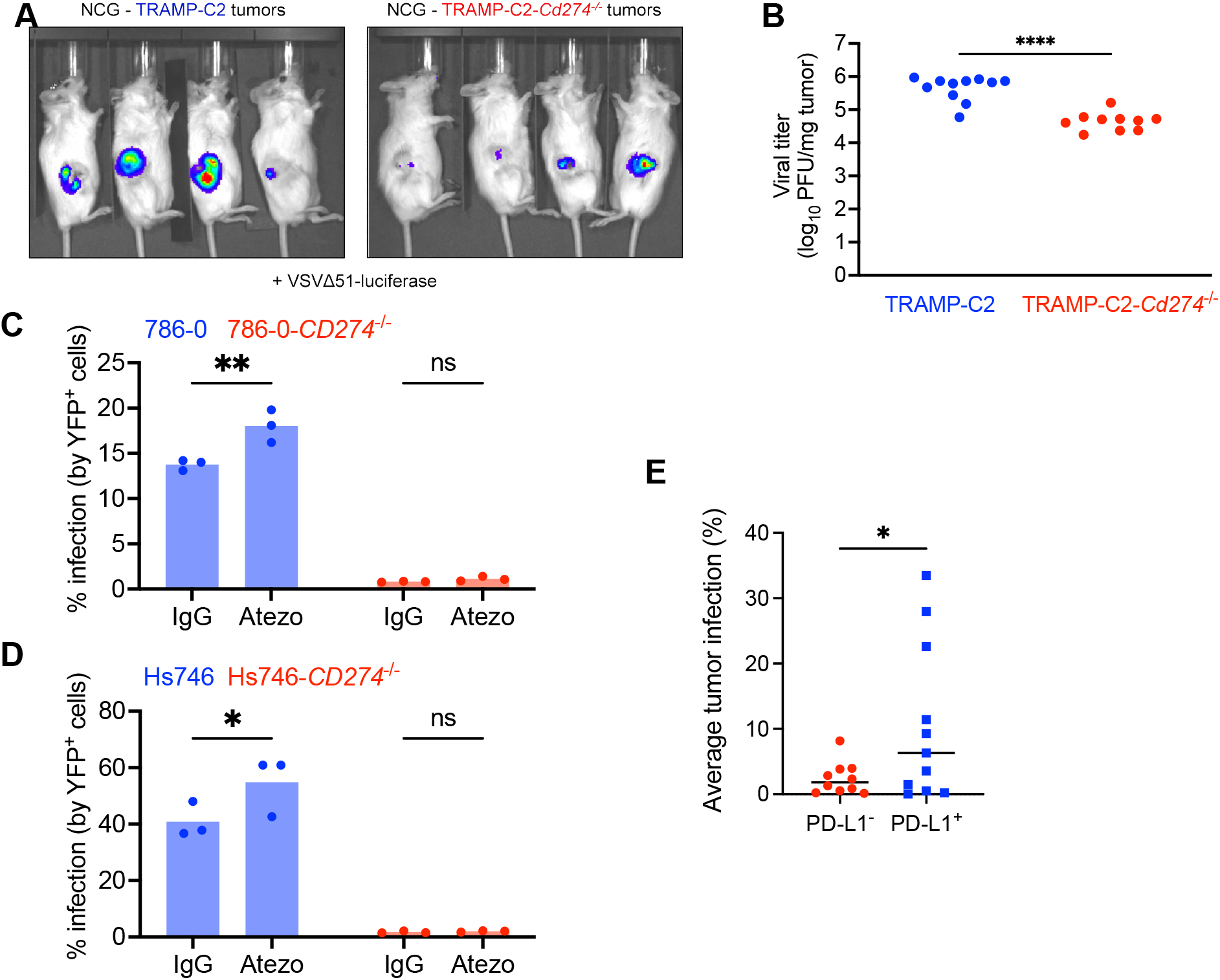
PD-L1 promotes OV infection in vivo. (**A**-**B**) Male NCG mice were implanted with subcutaneous TRAMP-C2 or TRAMP-C2-*Cd274*^-/-^ cells, and injected intra-tumoral with 10^8^ PFU of VSV1′51 expressing a luciferase reporter. 24 hours post-infection, mice were injected subjected to bioluminescence imaging, and tumors were homogenized to quantify viral titers by plaque assay. The bioluminescence images are representative of 2 experiments with similar results. Statistical analysis by unpaired two-tailed Student’s t-test. ****:p<0.0001. (**C-D**) 786-0 (C) and Hs746 (D) cells were treated with 5 μg of atezolizumab for 24 hours prior to VSV1′51-YFP infection at MOI 1 or 0.1, respectively. 24 hours post-infection, cells were analyzed by flow cytometry to quantify the viral YFP reporter. Experiments depicted are representative of 3 performed with similar results. Statistical analysis by two-way ANOVA with Šídák’s correction for multiple comparisons. (**E**) PD-L1 tumor status (where PD-L1^+^ tumors are defined as >1% PD-L1^+^ tumor/immune cells) plotted against average tumor infection (percentage of tumor explant area that is YFP^+^). Statistical analysis by two-tailed unpaired Student’s t-test. *:p<0.05

We next investigated if atezolizumab, the clinically approved PD-L1 antibody used in checkpoint inhibition immunotherapy, triggered PD-L1 function and enhanced OV infection. Pre-treatment of the 786-0 and Hs746 cells with atezolizumab significantly enhanced OV infection compared to the isotype control (Figures 7C-D); as expected, atezolizumab treatment had no impact on OV infection in PD-L1-deficient cells.

Lastly, we asked if PD-L1 favoured OV infection in primary human cancer reasoning that PD-L1^+^ tumors should have higher rates of infection, based on the totality of the data presented thus far. We obtained fresh tumor biopsies and subjected them to both *i)* ex vivo VSVΔ51-YFP infection for 48 hours (Figure S8A); and *ii)* PD-L1 immunohistochemistry to determine the PD-L1 status of tumors (PD-L1^+^ tumors are defined as ≥1% of tumor/immune cells staining for PD-L1, in accordance with clinical protocols) (Figure S8B). In a cohort of 21 patient tumors (Table S3), PD-L1^+^ tumor explants were significantly better infected compared to PD-L1^-^ explants (Figure 7E). Importantly, infection did not correlate with pre-infection tumor viability (Figure S9A) nor degree of biopsy necrosis (Figure S9B), suggesting that this analysis was not confounded by tissue viability, which was very high in most biopsies analyzed. Tumors derived from male and female patients were equally represented in terms of PD-L1 status (Figure S9C) and infection levels (Figure S9D). Therefore, in a cohort of tumors of diverse origins and treatment history, PD-L1 marked tumors that were more likely to be infected by OVs, corroborating our results in human tumors, and revealing an unappreciated role of PD-L1 as an OV infectivity biomarker.

## DISCUSSION

Here, we show that PD-L1 inhibits the type I IFN response and enhances OV infection via a pro-glycolytic shift in cancer cells resembling of Warburg metabolism. The requirement for engagement by an extracellular binding partner strongly suggests that this function is mediated by some signaling capacity of PD-L1. While the idea of PD-L1 “reverse signaling” has quickly gained traction there is still a lack of data towards understanding if PD-L1 functions requires cross-linking or conformational changes triggered by other proteins. Our data showing that CD80, a PD-1 fusion protein or monoclonal antibodies, including those clinically used, boost PD-L1 functions add to this debate suggesting that PD-L1 needs to be engaged to mediate its cell intrinsic functions.

PD-L1 has previously been shown to modulate type I IFNs with biochemically, transcriptomic and bioinformatic approaches. Surprisingly, some of these early studies show that PD-L1 inhibits type I IFN responses, while others showed the opposite (*23–25*). Overall, the mechanisms underlying this context-specificity of PD-L1 function are unknown. It is possible that PD-L1 signals differently depending on the cell type. What may be underpinning these signaling differences is the extensive glycosylation of PD-L1, which is responsible for approximately 50% of its observed molecular weight. Different cancer cell types/lines express different PD-L1 glycoforms (*54*), and differential glycosylation influences PD-L1 interactions (*55*), and, potentially, its downstream signaling. Our discovery that PD-L1 promotes glycolysis creates an intriguing link with PD-L1 glycosylation, which warrants further investigation.

Mechanistically, we found that PD-L1 regulates type I IFNs by promoting Warburg metabolism. Strengthening our finding, previous work suggested a link between PD-L1 expression and metabolism. For example, a correlation was found between tumor PD-L1 expression and PET signal in different cancer types (*56*), similar to what we observed in vivo. Additionally, PD-L1 was shown to influence aerobic glycolysis and other metabolic pathways in cancer cells (*20, 21, 57*). The biological consequences of PD-L1-mediated metabolic shifts remain understudied, particularly given the fact that PD-L1 is expressed on cells with metabolic functions, such as pancreatic islet cells (*1*), warranting more work investigating the impact of metabolic regulation by PD-L1. We have linked PD-L1 regulation of type I IFN to lactate produced during Warburg metabolism. Lactate is no longer considered simply a waste product, and is now known to be involved in the regulation of key oncogenes and tumor suppressor genes (*58*) as well as inflammation (*59*), and a novel post-translational modification (lactylation) involving addition of lactate to lysine and phenylalanine residues has been shown (*60, 61*).

Certainly, our data shed new light on the combination of PD-L1 blockade with OVs. The rationale behind this combination lies in the fact that OV infection leads to up-regulation of PD-L1 in many tumor models (*30, 35*), and therefore blocking PD-L1 will unleash the full range of anti-tumor immunity induced by OVs. Indeed, OVs combined with anti-PD-1/PD-L1 led to improvements in tumor immune infiltrate and the activation status of immune cells (*30, 62, 63*). At the same time, it is now reasonable to test if anti-PD-L1 triggers the ability of PD-L1 to enhance OV infection in tumors, independent of its effect on anti-tumor immunity. In accordance with our in vitro and in vivo data, in a small cohort of cancer patients we found that PD-L1 expression predicted susceptibility to OV infection with only one PD-L1^-^ tumor well-infected ex vivo. Since PD-L1 in tumors is regularly measured in clinical settings, it will be interesting to determine if this relationship between PD-L1 and OV infection holds in future trials.

Lastly, from a clinical perspective, the fact that PD-L1 antibodies can trigger PD-L1 activity is highly relevant. It is tempting to speculate that these novel, cell-intrinsic functions of PD-L1 are being modulated in tumors of patients undergoing anti-PD-L1 therapy, and this may contribute to anti-tumor efficacy (or lack thereof) of these therapeutic agents. Investigation in appropriate murine models of cancer is needed to elucidate the role of cell-intrinsic PD-L1 function on checkpoint blockade efficacy.

## MATERIALS AND METHODS

### Cell lines

Cell lines were cultured at 37°C in a humidified atmosphere containing 5% CO_2_. TRAMP-C2 were maintained in DMEM supplemented with 5% FBS, 5% NuSerum, 0.005 mg/mL bovine insulin, 10 nM dehydroisoandrosterone, 100 U/mL penicillin, 100 ug/mL streptomycin, 10 ug/mL gentamicin sulfate, and 20 mM HEPES. 786-0, Hs746, HEK293T, L929-ISRE, and Vero cells were maintained in DMEM supplemented with 10% FBS, 100 U/mL penicillin, 100 ug/mL streptomycin, 10 ug/mL gentamicin sulfate, and 20 mM HEPES. Cells were regularly tested for mycoplasma using PCR protocol adapted from (*64*). 786-0 and Vero cells were a gift from Dr. John Bell (OHRI), Hs746 were purchased from ATCC, L929-ISRE generated by Dr. Bruce Beutler (UT Southwestern) were a gift from Dr. Subash Sad (uOttawa), and HEK293T were a gift from Dr. Ian Lorimer (OHRI).

### Generation of cell line variants

Single-guide RNA (sgRNA) targeting exon 3 of the *Cd274* gene (sequence: GTATGGCAGCAACGTCACGA) was cloned into the Cas9-expressing lentiCRISPR v2 vector according to lentiCRISPR cloning protocol from the Zhang lab; lentiCRISPR v2 was a gift from Feng Zhang (Addgene plasmid 52961; http://n2t.net/addgene:52961; RRID: Addgene_52961). TRAMP-C2 were transiently transfected with plasmid, and subsequently treated with murine IFN-γ (Peprotech) to up-regulate PD-L1 in all cells, except those with deletion of PD-L1. PD-L1^-^ cells were isolated by FACS. To generate 786-0 and Hs746 *CD274*^-/-^, cells were electroporated with a ribonucleoprotein complex of ATTO550-labeled gRNA (IDT; sequences: UGG CUG CAC UAA UUG UCU AUG UUU UAG AGC UAU GCU; AUU UAC UGU CAC GGU UCC CAG UUU UAG AGC UAU GCU; AGC UAC UAU GCU GAA CCU UCG UUU UAG AGC UAU GCU; UUG AAG GAC CAG CUC UCC CUG UUU UAG AGC UAU GCU) and recombinant Cas9 (IDT) using protocols modified from the Alt-R CRISPR-Cas9 system (IDT), and subsequently treated with human IFN-γ (Peprotech) to up-regulate PD-L1 in all cells except those with deletion of PD-L1. PD-L1^-^ cells were isolated by FACS.

To stably express PD-L1 or control vector in TRAMP-C2-*Cd274*^-/-^, the cDNA encoding full-length murine PD-L1 was cloned into the retroviral vector pQCXIN-IRES-Thy1.1, and the resulting pQCXIN-PD-L1-IRES-Thy1.1 plasmid was transfected into HEK293T cells along with pCMV-VSV-G (a gift from Bob Weinberg; Addgene plasmid 8454; http://n2t.net/addgene:8454; RRID:Addgene_8454) and pCL-Eco (a gift from Inder Verma; Addgene plasmid 12371; http://n2t.net/addgene:12371; RRID:Addgene_12371) to generate retrovirus. TRAMP-C2-*Cd274*^-/-^ were infected with this retrovirus (or retrovirus encoding pQCXIN-IRES-Thy1.1 as empty vector control) supplemented with 8 µg/mL polybrene. Cells staining positively for PD-L1 and Thy1.1 (or Thy1.1 only for empty vector control) were isolated by FACS.

### Oncolytic virus production and infection

VSVΔ51-YFP, VSV WT, VSVΔ51-firefly luciferase, and vaccinia virus were gifts from Dr. John Bell (OHRI). The original virus stock was propagated on Vero cells (at MOI 0.01) and cell supernatant isolated 16-20 hours later for concentration of virus by high-speed centrifugation. All subsequent virus stocks were generated from the original stock to avoid genetic drift. VSVι151 titers were quantified by plaque assay using methods previously described (*65*).

TRAMP-C2, 786-0, and Hs746 were infected with VSVι151-YFP or VSV WT by first removing and washing out culture media with PBS and adding low volume of virus diluted in cold DMEM to MOIs ranging from 0.001 to 100. After incubation at 37°C + 5% CO_2_, supplemented growth media was added.

Alternatively, TRAMP-C2 were infected with GFP-expressing B19R^-^ vaccinia virus (Copenhagen strain) at MOI 0.1 for 48 hours.

### In vitro treatments

Cells were treated with the following reagents, at concentrations and times indicated in figure legends: recombinant murine IFN-β (PBL Assay Sciences), poly(I:C) (Invivogen), actinomycin D (Sigma), recombinant mouse PD-1-Fc chimeric protein (R&D Systems) or human IgG1 control (R&D Systems), afatinib (Selleck Chemicals), SB431542 (Selleck Chemicals), estradiol/E2 (Sigma), dihydrotestosterone/DHT (Sigma), sodium oxamate (Selleck Chemicals), (R)-GNE-140 (Selleck Chemicals), sodium lactate (Sigma), or oligomycin A (Selleck Chemicals).

### Coomassie staining

Culture media was removed and washed out with PBS, and cells were fixed with 3:1 methanol:acetic acid solution for 1-3 hours. Cells were then rinsed in tap water and stained with Coomassie Blue solution for 30 minutes and rinsed with tap water to remove excess dye.

### Flow cytometry

Cells were briefly trypsinized, washed, and resuspended in PBS for staining. The cell suspension was stained with Zombie NIR Fixable Viability Dye or Zombie Aqua Fixable Viability Dye (BioLegend) to label dead cells, followed by incubation with rat anti-mouse CD16/CD32 (clone 2.4G2, BD Biosciences) to block FcγRII/III receptors. Cells were then washed and incubated with fluorescently labeled antibodies. Cells were washed and resuspended in PBS and analyzed using an LSRFortessa (BD Biosciences) or Celesta (BD Biosciences), or isolated using MoFlo XDP (Beckman Coulter) or MA900 (SONY). Data were analyzed using FlowJo (Tree Star Inc.).

For experiment with murine cells, the following antibodies were used: BV421-anti-PD-L1 (BD, clone MIH5), PE-Cy5-anti-CD80 (BioLegend, clone 16-10A1), PE-Cy7-anti-PD-1 (BioLegend, clone 29F.1A12), PE-anti-LDLR (R&D Systems, clone 263123). For experiments with human cells, BV421-anti-PD-L1 (BD, clone MIH1) was used.

### RNA isolation, cDNA synthesis, and qPCR

RNA was isolated using GenElute RNA Miniprep Kit (Sigma) as per manufacturer’s protocol. cDNA was synthesized using iScript Reverse Transcription Supermix (Bio-Rad) as per manufacturer’s protocol. qPCR was run using iTaq Universal SYBR Green Supermix (Bio-Rad), using primers listed in the table below. qPCR data normalized to murine *Gapdh* gene or human *18S* gene using the 2-^ΔΔCt^ method.

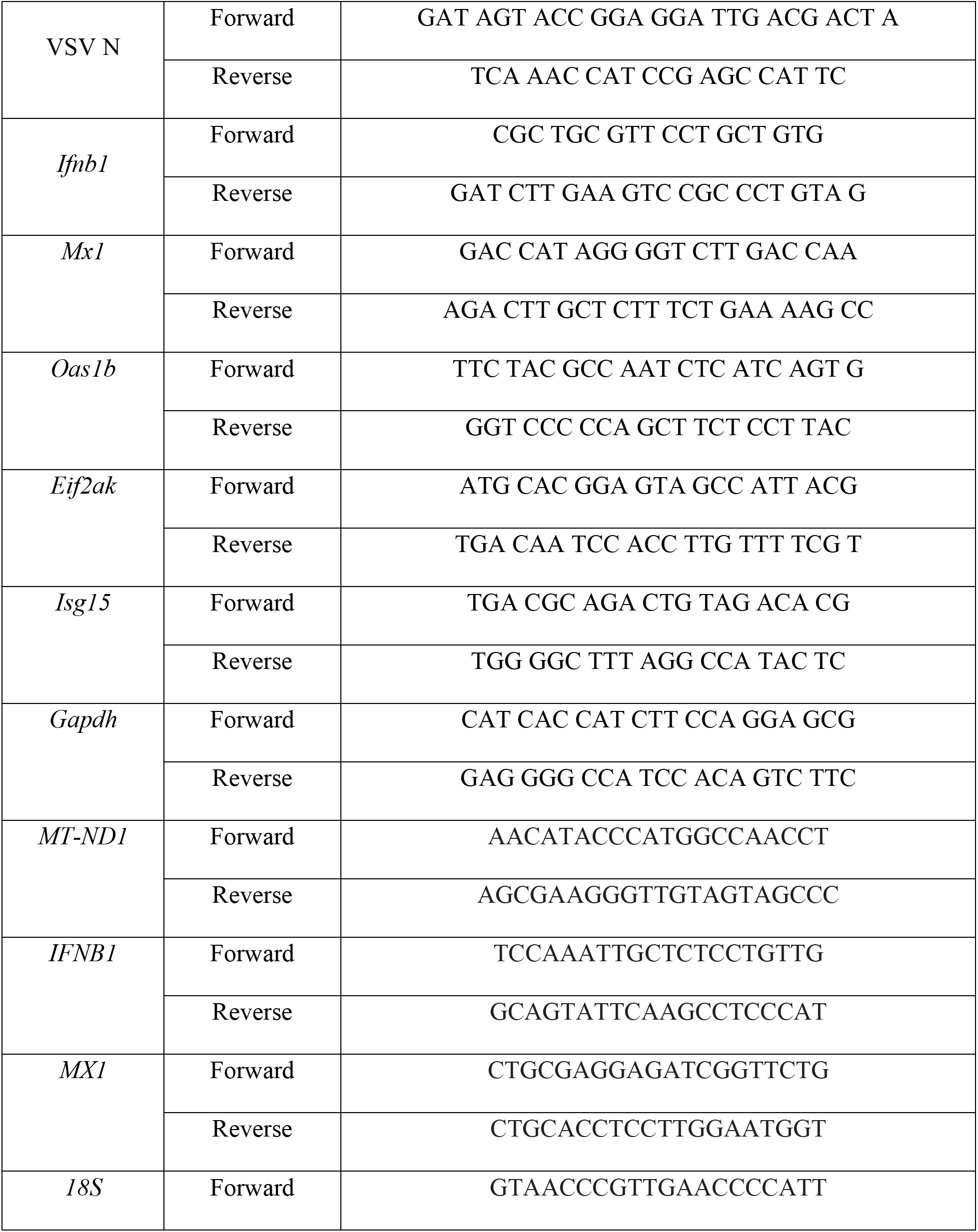
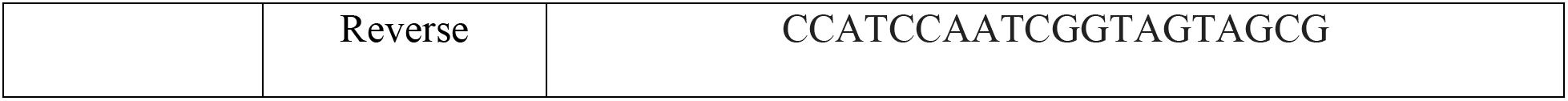

## ELISA

IFN-β was analyzed in culture supernatant following VSVΔ51-YFP/VSV WT infection or transfection with poly(I:C) (Invivogen), using Mouse IFN-β Quantikine ELISA Kit (R&D Systems), as per manufacturer’s instructions.

### Western blotting

Following infection or treatment with recombinant murine IFN-β (PBL Assay Sciences), cells were lysed in RIPA supplemented with cOmplete protease inhibitor cocktail (Roche) and PhosSTOP phosphatase inhibitor cocktail (Roche). Protein concentration was quantified by BCA Assay using MicroBCA Protein Assay Kit (ThermoFisher), and samples were denatured in Laemmli buffer (Bio-Rad) supplemented with 5% β-mercaptoethanol. Proteins were separated on 8-12% polyacrylamide (acrylamide/bis-acrylamide 37.5:1, Bio-Rad) gel at 60 mA, and transferred to a PVDF membrane for 90 minutes at 100V. Membranes were probed with the following primary antibodies diluted in 5% w/v BSA in 1X TBS + 0.1% Tween20: anti-pIRF3 S396 (CST), anti-IRF3 (CST), anti-pTBK1 S172 (CST), anti-TBK1 (CST), anti-pSTAT1 Y701 (CST), anti-STAT1 (CST), anti-pSTAT3 Y705 (CST), anti-STAT3 (CST), anti-PD-L1 (Abcam), anti-PD-1 (CST), and anti-β-actin (CST). Membranes were further probed with appropriate species-specific HRP-conjugated secondary antibodies (CST) and developed using ECL reagent (Bio-Rad).

### In vitro antibody treatment

Anti-IFNAR1 (clone MAR1-5A3) was purchased from Leinco. PD-L1 monoclonal antibody clones 6E11, 17H9, and 27C11 were gifts from Dr. Ira Mellman (Genentech). Tecentriq® (atezolizumab) was purchased from The Ottawa Hospital Pharmacy department. TRAMP-C2, 786-0, or Hs746 cells were pre-treated with these antibodies or appropriate isotype controls (Leinco) for 24 hours prior to further manipulation/analysis at concentrations indicated in figure legends.

### Type I interferon reporter assay

100 µL of cell culture supernatant was isolated from cultured cells and placed on adherent L929-ISRE cells (expressing luciferase under the control of a type I interferon sensitive ISRE promoter) for 4-6 hours. Luciferase expression was assessed using Luciferase Assay System (Promega) and luminescence measured on plate reader (BioTek).

### RNA-seq sample preparation

RNA was collected as described above from mock-infected TRAMP-C2 cells and TRAMP-C2-*Cd274*^-/-^ or those same cells infected with VSVΔ51-YFP at MOI 0.1 for 8 hours.

### RNA-seq library preparation and sequencing

Total RNA was quantified using a NanoDrop Spectrophotometer ND-1000 (NanoDrop Technologies, Inc.) and its integrity was assessed on a 2100 Bioanalyzer (Agilent Technologies). Libraries were generated from 250 ng of total RNA as follows: mRNA enrichment was performed using the NEBNext Poly(A) Magnetic Isolation Module (New England BioLabs). cDNA synthesis was performed using the NEBNext RNA First Strand Synthesis and NEBNext Ultra Directional RNA Second Strand Synthesis Modules (New England BioLabs). The remaining steps of library preparation were done using the NEBNext Ultra II DNA Library Prep Kit for Illumina (New England BioLabs). Adapters and PCR primers were purchased from New England BioLabs. Libraries were quantified using the Quant-iT™ PicoGreen^®^ dsDNA Assay Kit (Life Technologies) and the Kapa Illumina GA with Revised Primers-SYBR Fast Universal kit (Kapa Biosystems). Average size fragment was determined using a LabChip GX (PerkinElmer) instrument. Libraries were then sequenced on a NovaSeq6000 S4 200 cycle (2 x 100bp) flow cell to an approximate depth of 30M reads per sample.

### RNA-seq processing and differential expression

Transcript quantification for each sample was performed using Kallisto (v0.45.0) (*66*) with the GRCm38 transcriptome reference and the -b 50 bootstrap option. The R package Sleuth (v0.30.0) (*67*) was then used to construct general linear models for the log-transformed expression of each gene across experimental conditions. Wald’s test was used to test for differential expression between groups and the resultant *p*-values were adjusted to *q*-values using the Benjamini-Hochberg false discovery rate method.

### Signalling inference and gene set scoring

Relative activities of 14 signalling pathways were inferred using the R package PROGENy (v.1.9.6) (*68*), which provides a prebuilt regression model for signalling activity based on consistently responsive genes. Single-sample gene set scoring was performed using the R package singscore (v1.17.) (*69*), which computes independent scores for each sample from rank-based statics for each gene in the set.

### MitoTracker/MitoSOX

TRAMP-C2 cells were briefly trypsinized and re-suspended in PBS. Cells were counted and 500,000 cells were stained with 500 nM MitoTracker Deep Red FM (ThermoFisher) and 10 µM MitoSOX Red Mitochondrial Superoxide Indicator (ThermoFisher) for 10 minutes at 37°C. Dye was washed out with PBS and cells were analyzed by flow cytometry within 1 hour.

### Mitochondrial DNA and nuclear DNA quantification

Mitochondrial DNA and nuclear DNA were isolated using organic solvent extraction-based protocol adapted from (*70*). qPCR performed as described above using primers amplifying MT-ND1 (for mitochondrial DNA) and 18S rRNA (for nuclear DNA).

### Seahorse extracellular flux analysis

20,000 TRAMP-C2 cells/well were seeded into 96-well Seahorse plate one day prior to the Seahorse assay. On the day of the assay, cells were equilibrated for one hour in DMEM supplemented with 4 mM glutamine, 1 mM sodium pyruvate, 25 mM glucose, at pH 7.4. Oxygen consumption rate (OCR) and extracellular acidification rate (ECAR) were measured by monitoring dissolved oxygen and pH using the XF96 extracellular flux analyzer (Seahorse Bioscience) above the cell monolayer under basal conditions and following treatment with oligomycin (1 μM), FCCP (0.5 μM), and rotenone + antimycin A (1 μM each).

### Glucose uptake

In vitro glucose uptake was measured following 10 minutes of uptake using the Glucose Uptake-Glo Assay kit (Promega), as per manufacturer’s instructions.

### Metabolomics

Levels of metabolites from TRAMP-C2 cells and culture media following 48 hours of culture were quantified by liquid chromatography mass spectrometry (LC-MS). Sample temperature was maintained on ice or dry ice where possible, and all solvents were MS grade and pre-equilibrated to -20°C.

Cell pellets and media/supernatant were collected to a pre-chilled 2 mL tube containing 6 washed ceramic beads (1.4 mm) and 230 µl of methanol:water (1:1). Samples were vortexed 10s and cell lysis was done by beating for 60 s at 2000 rpm (bead beating was done twice) after adding 220 µL of acetonitrile. Samples were then incubated with a 2:1 dichloromethane:water solution on ice for 10 minutes. The polar and non-polar phases were separated by centrifugation at 4000g for 10 minutes at 1°C. The upper polar phase was dried using a refrigerated CentriVap Vacuum Concentrator at -4°C (LabConco Corporation, Kansas City, MO). Samples were resuspended in water and run on an Agilent 6470A tandem quadruple mass spectrometer equipped with a 1290 Infinity II ultra-high-performance LC (Agilent Technologies) using the Metabolomics Dynamic MRM Database and Method (Agilent), which uses an ion-pairing reverse phase chromatography. This method was further optimized for phosphate-containing metabolites with the addition of 5 µM InfinityLab deactivator (Agilent) to mobile phases A and B, which requires decreasing the backflush acetonitrile to 90%. Multiple reaction monitoring (MRM) transitions were optimized using authentic standards and quality control samples. Metabolites were quantified by integrating the area under the curve of each compound using external standard calibration curves with Mass Hunter Quant (Agilent). No corrections for ion suppression or enhancement were performed, as such, uncorrected metabolite concentrations are presented.

### Enzymatic measurement of L-lactate

L-lactate was quantified in culture supernatant using colorimetric assay (Abcam), as per manufacturer’s protocol.

### Mice and tumor injection

NCG mice (NOD-*Prkdc*^em26Cd52^*Il2rg*^em26Cd22^/NjuCrl, lacking functional T, B, and NK cells) were purchased from Charles River Laboratory and maintained at the University of Ottawa. For all experiments, male mice were used to match the male origin of injected prostate cancer cell lines (TRAMP-C2). To generate subcutaneous tumors, 1-2*10^6^ cancer cells were resuspended in Matrigel (BD Biosciences) and injected in the left flank.

### In vivo VSV1′51 infection and bioluminescence imaging

For in vivo treatments of NCG mice with VSV1′51-firefly luciferase, subcutaneous TRAMP-C2 tumors were injected for 24 hours with 10^8^ PFU of virus by intra-tumoral injection of 100 µL of a 10^9^ PFU/mL solution in sterile PBS. Mice were injected when tumors reached 750 mm^3^.

24 hours post-infection with VSV1′51-firefly luciferase, tumor-bearing mice were subjected to IVIS imaging. Mice were injected intraperitoneally with D-luciferin (Perkin Elmer) at a dose of 150 mg/kg for 15 minutes prior to isoflurane anesthesia. Mice were imaged using an IVIS Spectrum (Perkin Elmer). Investigators were not blinded during imaging.

### Ex vivo virion quantification of infected mouse tumors

Tumors were resected, weighed, and flash frozen in dry ice. Later, tumors were mechanically lysed using a TissueLyser II (Qiagen), in PBS supplemented with cOmplete protease inhibitor cocktail (Roche), using glass beads. Cell debris removed via centrifugation and 70 µm filters, and supernatant serially diluted for plaque assay as described above. Titers reported in PFU/mg of tumor.

### [^18^F]-FDG PET

Tumor-bearing mice (tumor diameter 750 mm^3^, n = 5-6 per group) fasted 5-8 hours before the imaging session were anesthetized with 2% isoflurane and intravenously injected with [^18^F]-FDG (5.98 ± 1.92 MBq) as a bolus over 30 sec via the lateral tail vein. Mice remained under isoflurane anesthesia in an induction chamber and body temperature was maintained with a heat lamp. Blood glucose measurements (mM) were taken via tail vein blood sampling before and after the PET scan using a MediSure Blood Glucose Monitoring System. 40 minutes after radiotracer delivery, mice were positioned in the PET scanner and a 10 min transmission scan was performed. A whole-body static scan was immediately acquired between 50-60 min using a Siemens DPET scanner. Emission data were corrected for attenuation and scatter, then reconstructed using the 3D-OSEM/MAP algorithm. Volumetric regions of interest (ROIs) were drawn conforming to tumor margins and quantified using a threshold of 25% of SUVmax (SUV_25_) (Savaikar et al., 2020). Uptake values obtained in Bq·cc^-1^ were converted to SUV using the injected dose (Bq) and animal body weight (kg).

### Bioinformatic analyses of human cell line RNA-seq

Analysis of expression levels of *CD274* transcript and HALLMARK_GLYCOLYSIS gene module (*71, 72*) in cell lines was performed using information from the RNA-Seq dataset E-MTAB-2706, which contains genome-wide transcriptome profiles of 675 cancer cell lines. Briefly, we downloaded the Reads Per Kilobase of transcript, per Million (RPKM) file containing all sequencing reads for each cell line, along with a file containing the gene names in various formats and a metadata file describing the samples. We then converted gene IDs in the expression matrix to gene symbols and removed duplicates and missing values. We next added sample names to the columns of the expression matrix, before leveraging the R-based package singscore (*69*) that allows for rank-based statistics to score a sample’s gene expression profile according to the activities of genes provided by curated gene modules. We set a cut-off on each axis at the mean of each value plus two standard deviations.

### Malignant cell PD-L1 expression and gene set activity

We have previously compiled and processed a collection of scRNA-seq data from 266 epithelial tumours (*46*). Automated annotation of cell types had been performed in conjunction with a copy number alteration inference to identify the malignant population of each data set. Only tumours with >200 malignant cells were retained in the cohort. Average PD-L1 expression (log-transformed counts per 10k transcripts) was calculated from this fraction. Gene set scores for the MSigDB Hallmark Hypoxia and Glycolysis gene sets were calculated for individual cells using the R package UCell, which implements a rank-based signature scoring method based on the Mann-Whitney U statistic.

### Immunohistochemistry

Fresh tumor biopsies were fixed in 10% neutral-buffered formalin for 24 hours prior to paraffin-embedding and sectioning. Sections were rehydrated using xylene and ethanol and subjected to antigen retrieval using 10 mM citrate buffer (pH 6.0) in a pressure cooker for 10 minutes. Sections were blocked in 10% normal goat serum (BioLynx) and incubated with anti-PD-L1 (clone 28-8, Abcam) overnight at 4°C. Endogenous peroxidase activity was quenched using 3% hydrogen peroxide and incubated with HRP-conjugated goat anti-rabbit (Cell Signaling Technologies). Detection performed using DAB Substrate (Vector Laboratories), followed by hematoxylin counterstaining. Sections were dehydrated and stabilized with mounting medium (ThermoFisher). PD-L1 expression was scored by a blinded trained pathologist. The percentage of tumor cells that stained with the PD-L1 antibody was estimated on each slide. The average intensity of staining was scored as 0 (no staining), 1+ (weak intensity), 2+ (moderate intensity), and 3+ (strong intensity). The cellular compartment with positive staining was noted; this included nuclear, cytoplasmic, or membranous. The background inflammatory cells were examined for positive staining as well, and percentage of necrotic cells.

### Ex vivo infection of patient tumors

Fresh tumor biopsies were cut into 2 x 2 x 2 cm cores using a biopsy punch and scalpel. Each tumor core placed into individual wells of 24-well plate with DMEM supplemented with 10% FBS, 100 U/mL penicillin, 100 ug/mL streptomycin, 10 ug/mL gentamicin sulfate, and 20 mM HEPES. Cores were infected in quadruplicate with MOI 100, 30, 3, 1, or 0.1 of VSV1′51-YFP (assuming ∼1*10^6^ cells per core) or uninfected control. Concurrently, tumor cores analyzed by alamarBlue Assay (ThermoFisher) as per manufacturer’s protocol. Tumors were infected for 48 hours prior to imaging YFP reporter from the virus using EVOS Imaging System (ThermoFisher). Virus infection was quantified by quantifying percentage of tumor area that is YFP^+^, using ImageJ (in a blinded fashion). The only exclusion criteria for a tumor in this study was lack of viability based on alamarBlue results.

### Statistical analyses

All in vitro experiments repeated at least 3 times, unless otherwise stated. Mouse studies were performed twice, unless otherwise stated. Statistical analyses were performed using GraphPad Prism (GraphPad). Experiments with two independent conditions were analysed by two-tailed unpaired Student’s t test, one-way ANOVA to compare three or more conditions, and two-way ANOVA (with Sidak’s correction for multiple comparisons) to compare groups influenced by two variables. Differences between experimental groups were considered significant when *p* < 0.05.

### Study approvals

Mouse studies were reviewed and approved by Animal Care and Veterinary Services at the University of Ottawa in accordance with the guidelines of the Canadian Institutes of Health Research. For human studies, informed and written consent in accordance with the Declaration of Helsinki was obtained from all patients, and approval was obtained from The Ottawa Hospital (REB 20180221-02H).

## LIST OF SUPPLEMENTARY MATERILAS

Figure S1 to S9

## Acknowledgements

We would like to thank: Dr. Tang and the uOttawa Flow Cytometry and Virometry Core Facility, Mr. Ortiz and the OHRI Flow Cytometry Core Facility, Dr. Naz and the uOttawa Metabolomics Core Facility (supported by the Terry Fox Foundation and uOttawa), Ms. Delic and the Global Tissue Consent program at The Ottawa Hospital, the Ottawa Methods Centre at OHRI, the uOttawa Pre-clinical Imaging Core (RRID:SCR_021832), the uOttawa Louise Pelletier Histology Core Facility (RRID: SCR_021737), and the ACVS staff. Thank you to all former and current Ardolino lab members for fruitful and enlightening discussions. We are also thankful to Dr. Roy for honest and insightful comments on the manuscript.

## Author Contributions

Conceptualization: J.J.H., M-C.B-D., and M.A.; Formal analysis: J.J.H., J.A-H., D.P.C., E.Y., A.B., S.F., M.P., C.F-M., M.F.C., B.H.R, and M-C.B-D; Funding acquisition: J.C.B., M.E.H., B.H.R., R.C.A., B.C.V., L.A.S., M-C.B-D, and M.A.; Investigation: J.J.H., J.A-H., A.H., D.P.C., C.T.D., M.M., E.Y., A.B., S.F., C.F-M., M.F.C., and M.P.; Methodology: J.J.H, B.H.R., R.C.A., M.E.H., and M-C.B-D; Resources: J.C.B., M.E.H., B.H.R., R.C.A., B.C.V., L.A.S., and M-C.B-D; Supervision: J.C.B., M.E.H., B.H.R., R.C.A., B.C.V., L.A.S., M-C.B-D, and M.A.; Visualization: J.J.H and M.A.; Writing-original draft: J.J.H. and M.A.; Writing-review and editing: all authors.

## Data availability

The transcriptomic data that support the findings of this study are openly available in GEO at https://www.ncbi.nlm.nih.gov/geo/query/acc.cgi?acc=GSE210884, accession number GSE210884. The metabolomic data that support the findings of this study will be deposited in the National Metabolomics Data Repository (NMDR).

