## Supplementary files for "More than a ligand: PD-L1 promotes oncolytic virus infection via a metabolic shift that inhibits the type I interferon pathway"

**SUPPLEMENTARY MATERIAL**

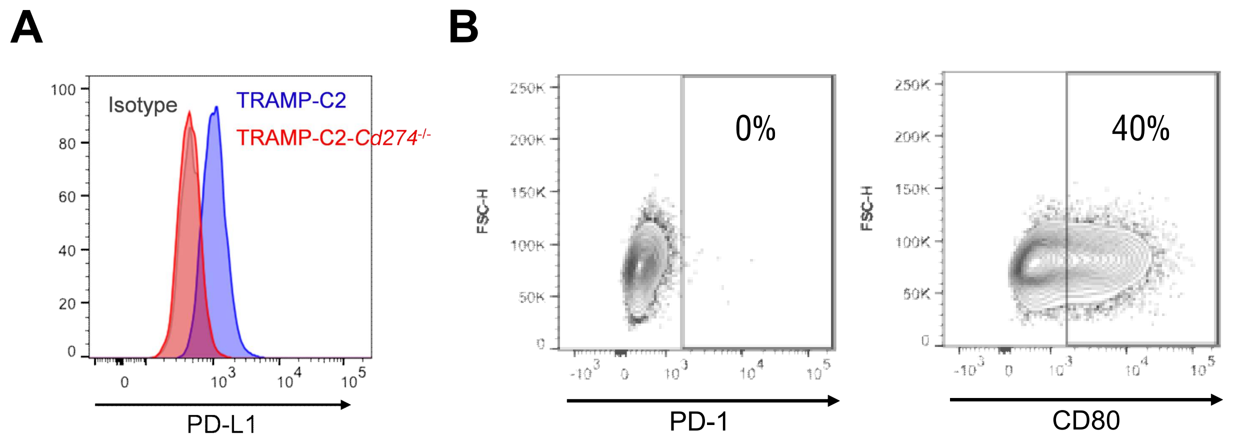


**FIGURE S1**

**Figure S1: PD-L1 engagement promotes oncolytic virus infection.** (**A**) TRAMP-C2 cells (blue) were transfected with Cas9 and gRNA targeting PD-L1 to generate TRAMP-C2-*Cd274*$^{-/-}$ (red). A representative plot depicting PD-L1 expression is shown. (**B**) Expression of PD-1 and CD80 in TRAMP-C2 cells. Representative flow plots are shown.

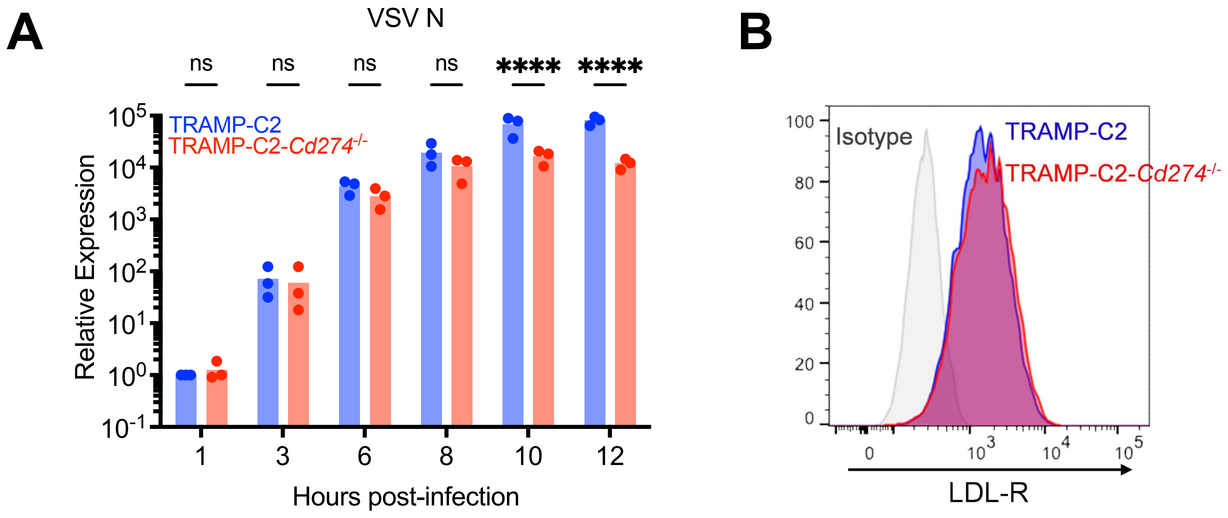


**FIGURE S2**

**Figure S2: PD-L1 does not regulate viral entry.** (**A**) TRAMP-C2 or TRAMP-C2-*Cd274*$^{-/-}$ were infected with VSVΔ51-YFP at MOI 0.1, and RNA collected at indicated times post-infection for qPCR analysis of the viral nucleocapsid (N) transcript. n=3 biological replicates depicted. Statistical analysis by two-way ANOVA with Šídák's correction for multiple comparisons. (**B**) Expression of the VSV entry receptor LDL-R in TRAMP-C2 (blue) or TRAMP-C2-*Cd274*$^{-/-}$ (red) cells. Representative histograms are shown.

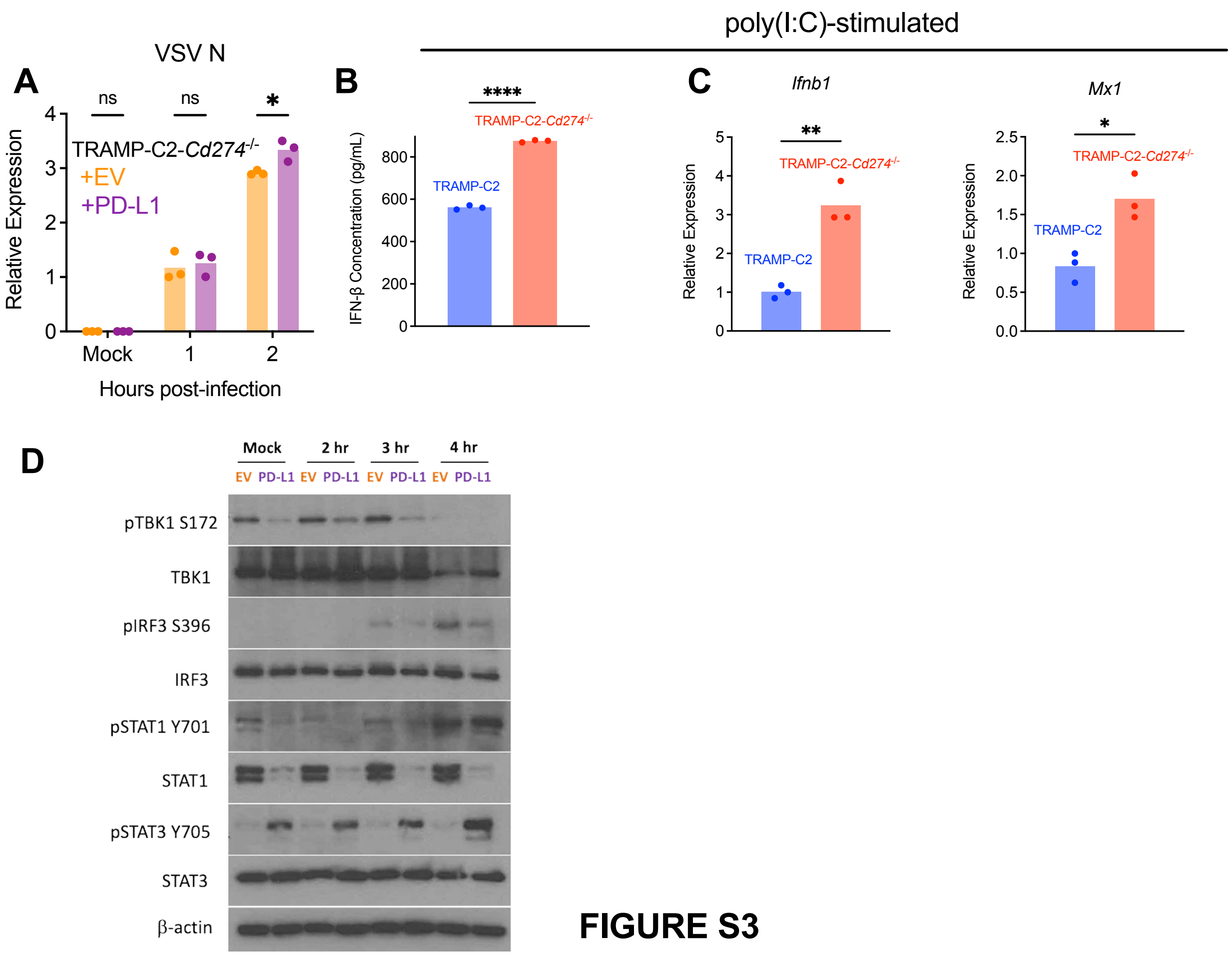


**FIGURE S3**

**Figure S3**: **PD-L1 inhibits the type I interferon response to oncolytic viruses.** (**A**) TRAMP-C2-*Cd274*$^{-/-}$ cells transduced with PD-L1 or empty vector were infected with VSVΔ51-YFP at MOI 0.1 and analyzed by qPCR at indicated times post-infection to quantify viral N transcripts. The data depicted are representative of 3 performed with similar results. Statistical analysis by two-way ANOVA with Šídák's correction for multiple comparisons. (**B** and **C**) Cells were transfected with 1 ug of poly(I:C) for 8 hours prior to quantification of IFN-β (by ELISA, B) and IFN-β and MX1 transcripts (by qPCR, C). The experiments depicted are representative of 3 performed with similar results. Statistical analysis by two-tailed unpaired Student's t-test. (**D**) TRAMP-C2-*Cd274*$^{-/-}$ cells transduced with PD-L1 or empty vector were infected with

VSVΔ51-YFP at MOI 0.1 and analyzed by western blotting at indicated times post-infection. The images depicted are representative of 3 performed with similar results.

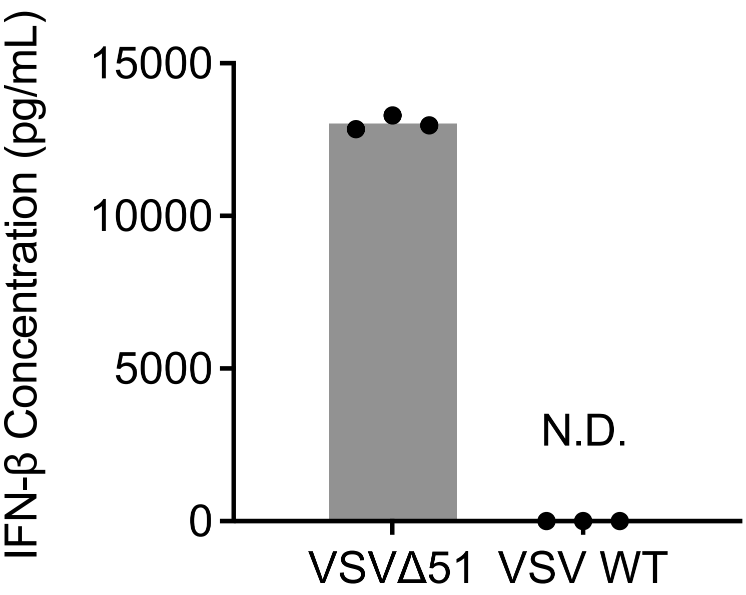


**FIGURE S4**

**Figure S4: VSV WT efficiently blocks type I IFN responses.** TRAMP-C2 cells were infected with VSVΔ51-YFP or VSV WT at MOI 0.1 for 12 hours prior to analysis by ELISA to quantify IFN-β in supernatant. The experiment depicted is representative of 2 performed with similar results. N.D. = not detected.

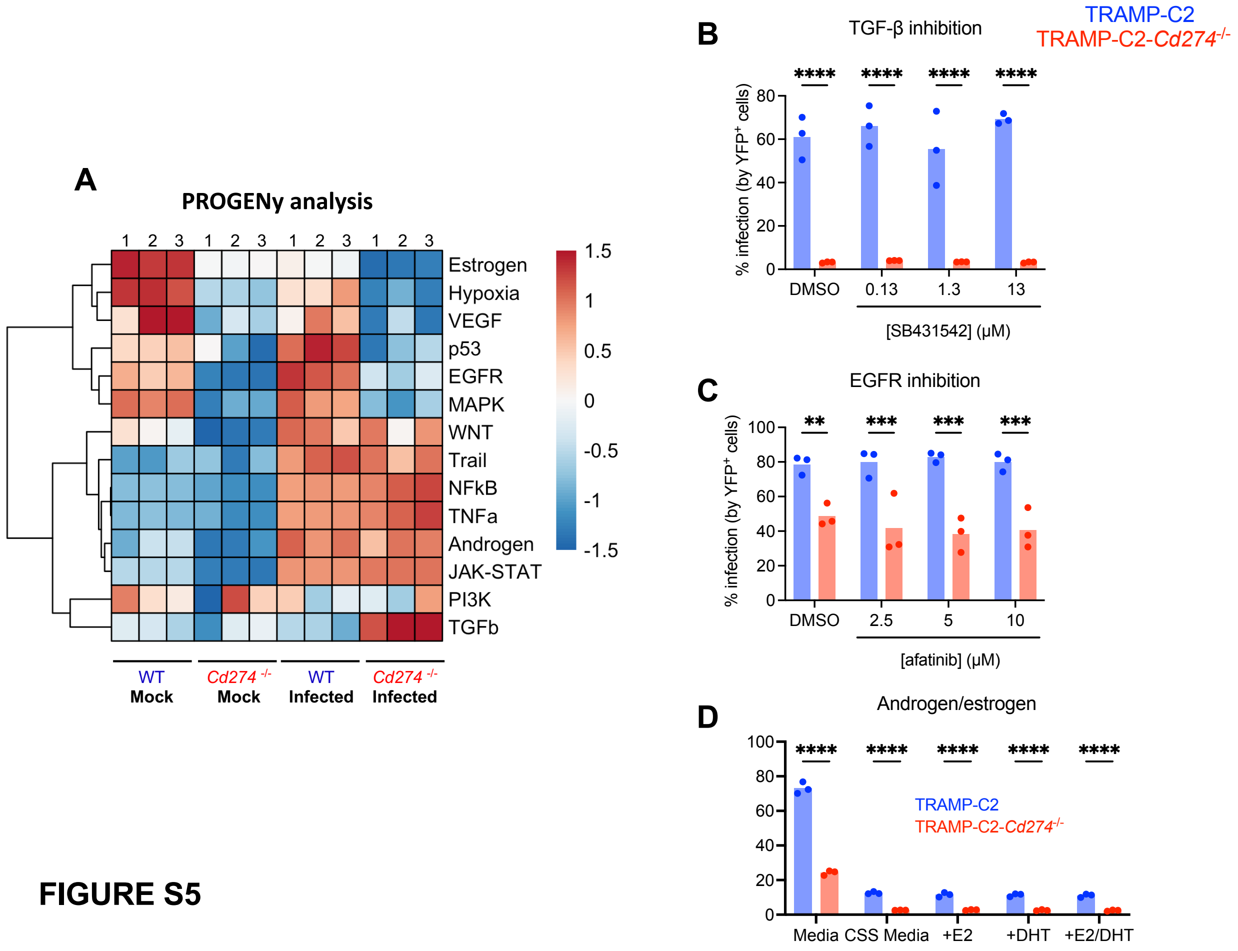


**FIGURE S5**

**Figure S5: TGF-β, EGFR, and androgen/estrogen pathways are not involved in PD-L1's promotion of viral infection.** (**A**) PROGENy pathway analysis of bulk RNA-seq performed on TRAMP-C2 and TRAMP-C2-*Cd274*$^{-/-}$ either mock-infected or infected with VSVΔ51-YFP at MOI 0.1 for 8 hours (3 biological replicates per condition). (**B-C**) TRAMP-C2 and TRAMP-C2-*Cd274*$^{-/-}$ cells were pre-treated with the TGF-βRI inhibitor SB431452 for 24 hours (**B**) or the EGFR inhibitor afatinib for 6 hours (**C**) at indicated concentrations (or DMSO as vehicle control), followed by infection with VSVΔ51-YFP at MOI 0.1 for 24 hours prior to analysis by flow cytometry for viral YFP reporter expression. Experiments depicted are representative of 2 with similar results. Statistical analysis by two-way ANOVA with Šídák's correction for multiple comparisons. (**D**) TRAMP-C2 and TRAMP-C2-*Cd274*$^{-/-}$ cells were cultured in phenol red-free

DMEM supplemented with 10% charcoal-stripped serum (CSS) for 48 hours prior to treatment with estradiol (E2) or dihydrotestosterone (DHT) for 24 hours. Following that, cells were infected with VSVΔ51-YFP at MOI 0.1 for 24 hours prior to analysis by flow cytometry for viral YFP reporter expression. Experiments depicted are representative of 2 with similar results. Statistical analysis by two-way ANOVA with Šídák's correction for multiple comparisons.

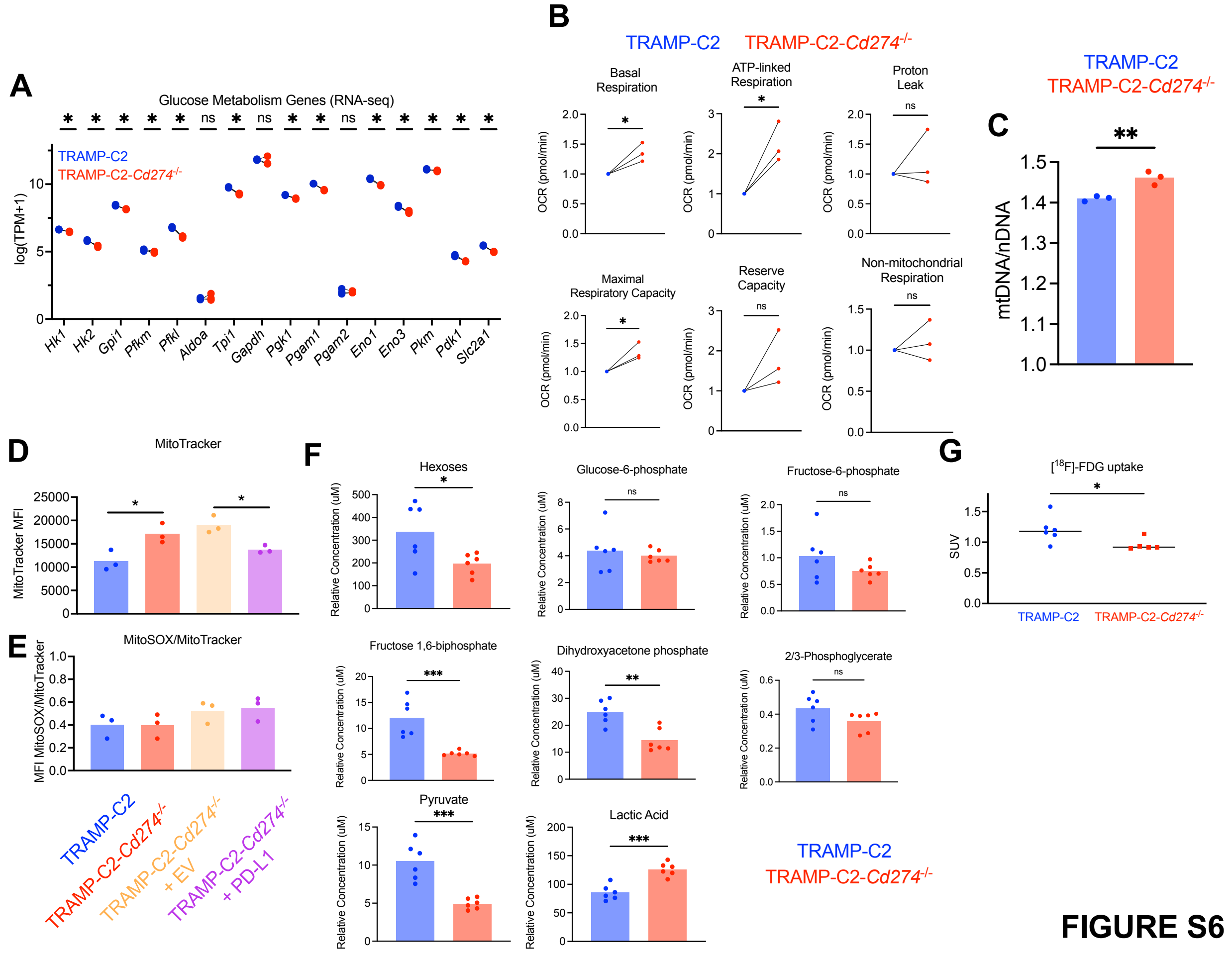


**Figure S6: PD-L1 promotes Warburg metabolism in cancer cells.** (**A**) Transcripts per million (TPM) for various glycolysis and glucose metabolism enzymes/transporters, from bulk RNA-seq experiment. **(B)** Cellular respiration parameters calculated using Seahorse mitochondrial stress test data. N=3 biological replicates are shown. For each replicate, data normalized to TRAMP-C2 values. Statistical analysis by two-tailed unpaired Student's t-test. (**C**) Relative quantification of mitochondrial DNA/mtDNA by qPCR (normalized to nuclear DNA/nDNA in each sample), by qPCR analysis. N=3 biological replicates. Statistical analysis by two-tailed unpaired Student's t-test. (**D**) Mean fluorescence intensity (MFI) of MitoTracker and MitoSOX dyes by flow cytometry for TRAMP-C2, TRAMP-C2-*Cd274*$^{-/-}$, TRAMP-C2-*Cd274*$^{-/-}$ + EV, and TRAMP-C2-*Cd274*$^{-/-}$ +

PD-L1 cells. MitoSOX MFI was corrected for total mitochondrial content by normalizing to MitoTracker MFI. Statistical analysis by one-way ANOVA with Šídák's correction for multiple comparisons. (**E**) Relative abundance of glycolysis metabolites in TRAMP-C2 and TRAMP-C2-*Cd274*$^{-/-}$ cells, measured in the untargeted metabolomics study. 6 biological replicates per cell line. Statistical analysis by unpaired two-tailed Student’s t-test. (**F**) Male NCG mice were implanted with subcutaneous TRAMP-C2 or TRAMP-C2-*Cd274*$^{-/-}$ tumors. Standardized uptake value (SUV) of $[^{18}F]$-fluorodeoxyglucose was assessed by PET imaging. Statistical analysis by two-tailed unpaired Student’s t-Test. *: $p<0.05$.

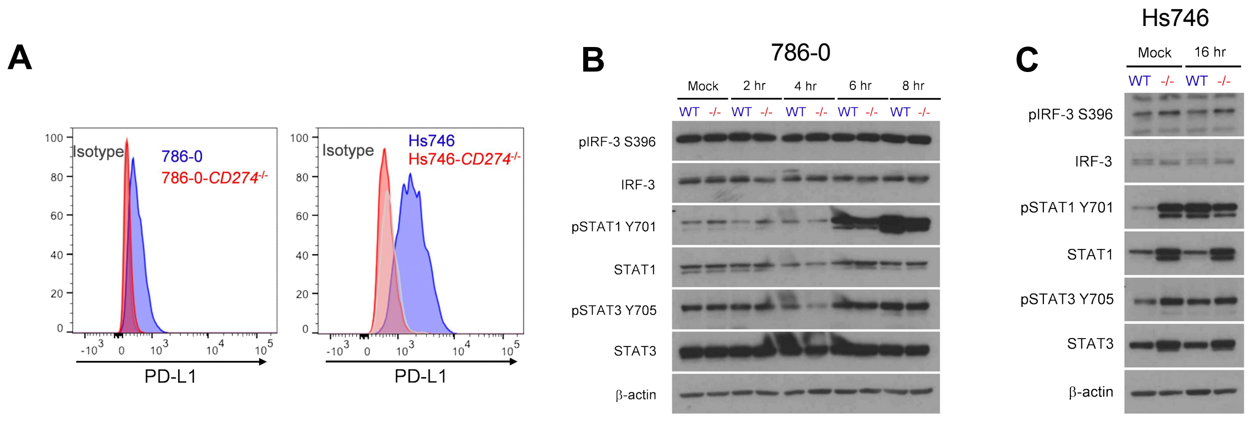


**FIGURE S7**

**Figure S7: PD-L1 alters type I interferon responses in human cancer cells .** 786-0 (left) and Hs746 (right) cells were electroporated with Cas9 and gRNA targeting PD-L1 to generate *CD274*$^{-/-}$ cells. Representative plots depicting PD-L1 expression are shown. (**B**) 786-0 and 786-0-*CD274*$^{-/-}$ cells infected with VSVΔ51-YFP at MOI 1 (or mock-infected) and analyzed by western blotting at indicated times post-infection. Images depicted are representative of 3 with similar results. (**C**) Hs746 and Hs746-*CD274*$^{-/-}$ cells infected with VSVΔ51-YFP at MOI 0.1 (or mock-infected) for 16 hours prior to western blotting analysis. Images depicted are representative of 3 with similar results.

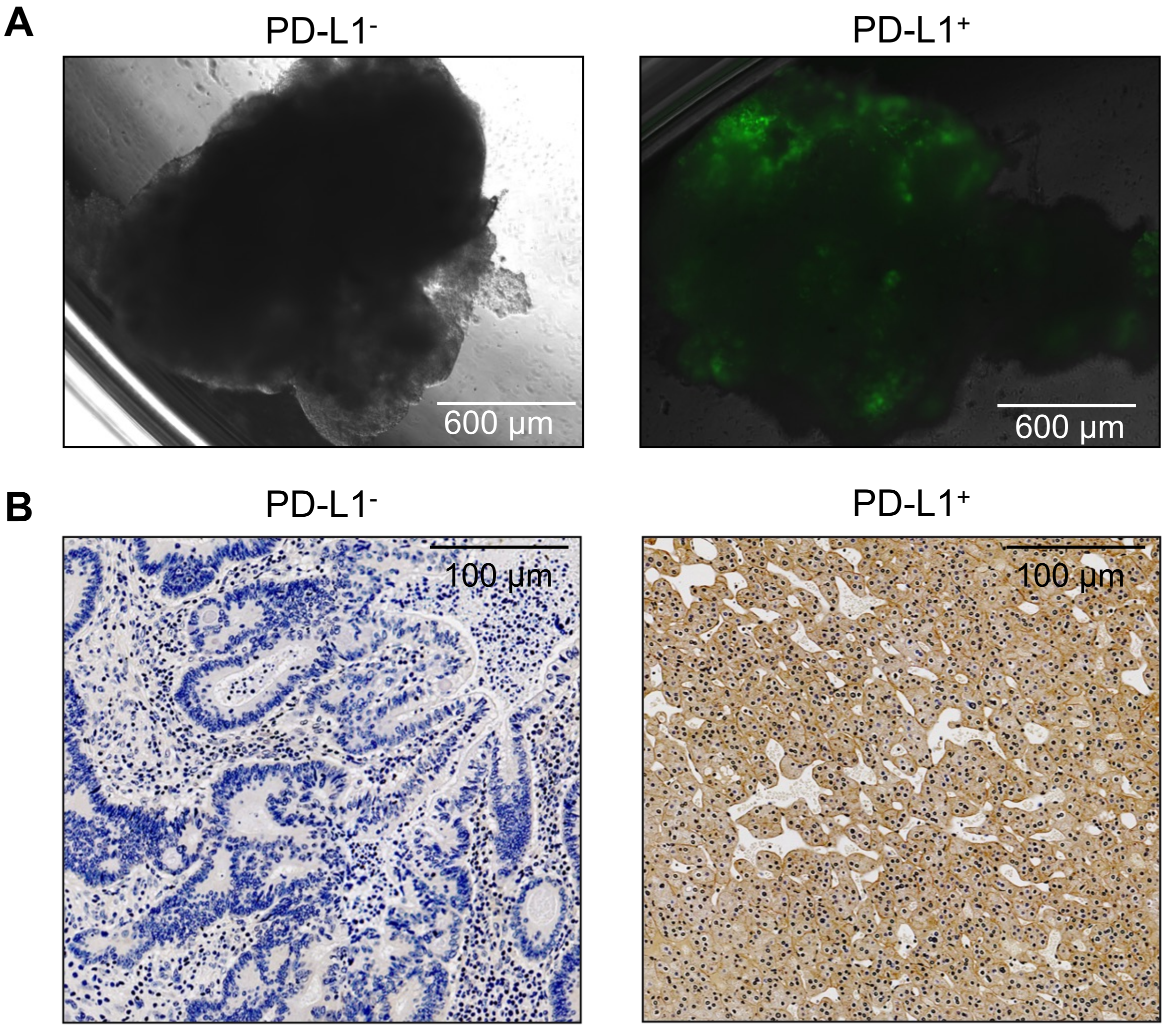


**FIGURE S8**

**Figure S8: assessment of PD-L1 by immunohistochemistry and oncolytic virus infection by fluorescent microscopy.** (**A**) Fluorescent imaging of tumor explant cores from patient biopsies infected with VSVΔ51-YFP for 48 hours prior to imaging for viral YFP reporter. Images of poorly infected (left) and well infected (right) tumors are depicted. (**B**) Patient tumor biopsies subjected to PD-L1 IHC to determine PD-L1 status of tumors. Images of PD-L1$^{-}$ (left) and PD-L1$^{+}$ (right) tumors are depicted.

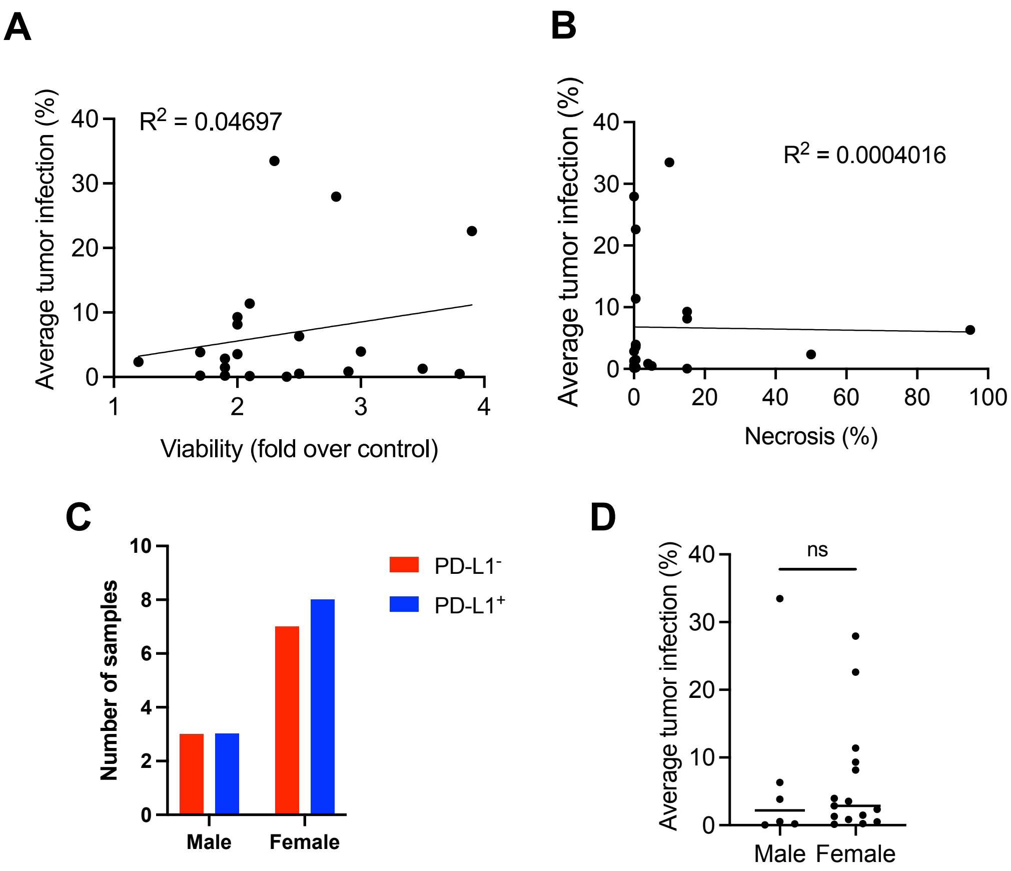


FIGURE S9

**Figure S9: PD-L1 is a biomarker of patient tumors responsive to oncolytic virus.** (**A-B**) Average tumor infection plotted against viability (assessed by alamarBlue) and necrosis (% of tumor section necrotic, assessed by histology). (**C-D**) Distribution of PD-L1$^+$ and PD-L1$^-$ tumors, and average tumor infection, between male and female donors. Statistical analysis by two-tailed unpaired Student's t-test.

**Table S1**. Metadata for tumor scRNA-seq data analyzed for PD-L1 and glycolysis gene set score.

| Cancer | Source | Accession | Platform | Sorted? | Patient # (>100 malignant cells) | Total cancer cell # | % mito threshold |
|---|---|---|---|---|---|---|---|
| Breast | Wu *et al.* (37) | ENA accession PRJEB35405 | 10x Genomics Chromium (3' v2) | NA | 4 | 4452 | 20% |
| Breast | Qian *et al.* (28) | http://blueprint.lambrechtslab.org | 10x Genomics Chromium (3') | NA | 10 | 8766 | 20% |
| Breast | Bassez *et al.* (38) | http://biokey.lambrechtslab.org/ | 10x Genomics Chromium (5') | NA | 62 | 38,765 | 15% |
| Colorectal | Lee *et al.* (26) | GEO Accession GSE144735 & GSE132465 | 10x Genomics Chromium (3' v2) | NA | 25 | 18,058 | 20% |
| Colorectal | Uhlitz *et al.* (27) | Direct from authors | 10x Genomics Chromium (3' v3) | NA | 8 | 2659 | 20% |
| Colorectal | Qian *et al.* (28) | http://blueprint.lambrechtslab.org | 10x Genomics Chromium (3') | NA | 11 | 8766 | 25% |
| Gastric | Sathe *et al.* (29) | https://dna-discovery.stanford.edu | 10x Genomics Chromium (3' v2) | NA | 7 | 6909 | 20% |
| Lung | Kim *et al.* (32) | GEO Accession GSE131907 | 10x Genomics Chromium (3' v2) | NA | 20 | 15,396 | 20% |
| Lung | Lambrechts *et al.* (31) | ArrayExpress Accessions E-MTAB-6149 & E-MTAB-6653 | 10x Genomics Chromium (3' v1/v2) | NA | 9 | 8036 | 20% |
| Lung | Qian *et al.* (28) | http://blueprint.lambrechtslab.org | 10x Genomics Chromium (3') | NA | 24 | 6794 | 20% |
| Lung | Laughney *et al.* (24) | GEO Accession GSE123904 | 10x Genomics Chromium (3' v2) | Viability (scatter & DAPI) | 8 | 3097 | 20% |
| Lung | Wu *et al.* (33) | GEO Accession GSE148071 | GEXSCOPE (Singleron Biotechnologies) | NA | 35 | 54,052 | 30% |
| Nasopharyngeal | Chen *et al.* (34) | GEO Accession GSE150430 | 10x Genomics Chromium (3' v2) | NA | 9 | 7400 | 20% |
| Ovarian | Geistlinger *et al.* (33) | GEO Accession GSE154600 | 10x Genomics Chromium (3' v2) | NA | 5 | 7479 | 20% |
| Ovarian | Qian *et al.* (28) | http://blueprint.lambrechtslab.org | 10x Genomics Chromium (3') | NA | 5 | 4967 | 20% |
| Pancreatic | Steele *et al.* (36) | GEO Accession GSE155698 | 10x Genomics Chromium (3') | NA | 15 | 10,495 | 20% |
| Pancreatic | Peng *et al.* (54) | GSA: CRA001160 | 10x Genomics Chromium (3' v2) | | 24 | 41,986 (all cell types) | 10% |
| Squamous cell carcinoma | Ji *et al.* (20) | GEO Accession GSE144236 | 10x Genomics Chromium (3' v2) | NA | 9 | 12,154 | 10% |

**Table S1**: Metadata for tumour scRNA-seq data analyzed for PD-L1 expression and glycolysis gene set score.

| Name | TRAMP-C2 | | | | | | TRAMP-C2-*Cd274*$^{-/-}$ | | | | | | Average | | Standard Deviation | |
|---|---|---|---|---|---|---|---|---|---|---|---|---|---|---|---|---|
| | 1 | 2 | 3 | 4 | 5 | 6 | 1 | 2 | 3 | 4 | 5 | 6 | TRAMP-C2 | TRAMP-C2-*Cd274*$^{-/-}$ | TRAMP-C2 | TRAMP-C2-*Cd274*$^{-/-}$ |
| 2/3-Phosphoglyceric acid | 0.440 | 0.310 | 0.531 | 0.477 | 0.490 | 0.361 | 0.274 | 0.390 | 0.288 | 0.393 | 0.402 | 0.395 | 0.435 | 0.357 | 0.084 | 0.059 |
| 2-Aminoadipic acid | 0.276 | 0.293 | 0.342 | 0.308 | 0.271 | 0.325 | 0.237 | 0.242 | 0.232 | 0.311 | 0.278 | 0.279 | 0.303 | 0.263 | 0.028 | 0.031 |
| 2-Deoxycytidine 5-monophosphate | 0.017 | 0.018 | 0.015 | 0.018 | 0.017 | 0.011 | 0.012 | 0.010 | 0.010 | 0.016 | 0.012 | 0.012 | 0.016 | 0.012 | 0.003 | 0.002 |
| 2-Deoxyribose 5-phosphate | 0.014 | 0.015 | 0.021 | 0.016 | 0.013 | 0.019 | 0.015 | 0.015 | 0.013 | 0.014 | 0.015 | 0.016 | 0.016 | 0.015 | 0.003 | 0.001 |
| Acetyl-CoA | 0.183 | 0.196 | 0.227 | 0.147 | 0.176 | 0.192 | 0.165 | 0.175 | 0.141 | 0.149 | 0.172 | 0.207 | 0.187 | 0.168 | 0.026 | 0.023 |
| Adenosine 3-5-cyclic monophosphate | 0.017 | 0.016 | 0.016 | 0.021 | 0.018 | 0.022 | 0.011 | 0.010 | 0.008 | 0.016 | 0.016 | 0.013 | 0.018 | 0.012 | 0.003 | 0.003 |
| Adenosine 5-diphosphate | 26.909 | 27.209 | 35.117 | 32.326 | 26.903 | 33.526 | 42.638 | 40.394 | 37.369 | 39.759 | 38.675 | 35.251 | 30.332 | 39.014 | 3.750 | 2.553 |
| Adenosine 5-monophosphate | 82.128 | 65.475 | 79.099 | 91.645 | 75.332 | 89.808 | 115.292 | 100.263 | 95.158 | 123.168 | 111.312 | 92.641 | 80.581 | 106.306 | 9.674 | 12.146 |
| Adenosine 5-triphosphate | 31.977 | 36.849 | 46.080 | 42.885 | 38.200 | 35.408 | 55.566 | 47.266 | 39.334 | 32.765 | 43.702 | 42.470 | 38.567 | 43.517 | 5.130 | 7.661 |
| Alanine/Sarcosine | 219.572 | 188.941 | 299.765 | 254.535 | 210.212 | 275.917 | 328.564 | 307.018 | 326.779 | 244.540 | 215.688 | 265.791 | 241.490 | 281.397 | 42.359 | 46.606 |
| alpha-D-Glucose-1-phosphate | 0.022 | 0.022 | 0.044 | 0.016 | 0.037 | 0.047 | 0.129 | 0.102 | 0.027 | 0.101 | 0.094 | 0.129 | 0.031 | 0.097 | 0.013 | 0.037 |
| alpha-Ketoglutaric acid | 0.572 | 0.949 | 0.932 | 0.764 | 0.805 | 0.640 | 0.511 | 0.470 | 0.514 | 0.560 | 0.522 | 0.575 | 0.777 | 0.525 | 0.152 | 0.038 |
| Asparagine | 0.091 | 0.065 | 0.091 | 0.095 | 0.055 | 0.131 | 0.153 | 0.191 | 0.147 | 0.145 | 0.175 | 0.133 | 0.088 | 0.157 | 0.027 | 0.022 |
| Aspartic Acid | 11.151 | 10.696 | 11.971 | 12.237 | 10.977 | 11.709 | 6.741 | 6.646 | 6.323 | 6.819 | 6.968 | 6.314 | 11.457 | 6.635 | 0.606 | 0.267 |
| beta-Nicotinamide adenine dinucleotide | 6.241 | 6.579 | 8.885 | 7.614 | 6.456 | 7.695 | 10.035 | 9.836 | 8.521 | 9.276 | 10.131 | 8.924 | 7.245 | 9.454 | 1.010 | 0.652 |
| cis-Aconitic acid | 0.117 | 0.112 | 0.113 | 0.144 | 0.138 | 0.118 | 0.111 | 0.104 | 0.099 | 0.091 | 0.063 | 0.100 | 0.124 | 0.095 | 0.014 | 0.017 |
| Citramalic acid | 0.121 | 0.195 | 0.195 | 0.180 | 0.201 | 0.107 | 0.147 | 0.128 | 0.139 | 0.222 | 0.167 | 0.175 | 0.166 | 0.163 | 0.042 | 0.034 |
| Citric acid / Isocitric acid | 9.129 | 9.081 | 10.670 | 10.306 | 10.673 | 14.036 | 9.353 | 10.862 | 7.783 | 23.525 | 7.862 | 8.851 | 10.649 | 11.373 | 1.809 | 6.059 |
| Citrulline | 0.809 | 0.719 | 0.869 | 0.859 | 0.652 | 1.074 | 1.740 | 1.842 | 1.671 | 1.627 | 1.759 | 1.579 | 0.830 | 1.703 | 0.146 | 0.096 |
| Coenzyme A | 0.275 | 0.278 | 0.318 | 0.314 | 0.317 | 0.257 | 0.311 | 0.272 | 0.290 | 0.305 | 0.328 | 0.291 | 0.293 | 0.300 | 0.026 | 0.020 |
| Creatine | 339.123 | 244.164 | 435.814 | 399.445 | 331.218 | 386.367 | 458.768 | 335.085 | 358.605 | 259.611 | 277.967 | 295.368 | 356.022 | 330.901 | 67.211 | 72.501 |
| Creatinine | 0.267 | 0.195 | 0.279 | 0.328 | 0.295 | 0.360 | 0.285 | 0.316 | 0.202 | 0.236 | 0.242 | 0.238 | 0.287 | 0.253 | 0.057 | 0.041 |
| Cystine | 0.003 | 0.001 | 0.003 | 0.004 | 0.003 | 0.007 | 0.010 | 0.012 | 0.007 | 0.006 | 0.006 | 0.007 | 0.003 | 0.008 | 0.002 | 0.002 |
| Cytidine | 0.081 | 0.106 | 0.092 | 0.122 | 0.098 | 0.101 | 0.074 | 0.076 | 0.080 | 0.087 | 0.057 | 0.072 | 0.100 | 0.074 | 0.014 | 0.010 |
| Cytidine 5-diphosphate | 7.740 | 7.128 | 10.026 | 10.031 | 8.744 | 10.206 | 12.701 | 11.569 | 11.140 | 10.906 | 11.276 | 10.093 | 8.979 | 11.281 | 1.321 | 0.857 |
| Cytidine 5-monophosphate | 1.677 | 1.716 | 1.758 | 2.217 | 1.673 | 1.609 | 2.012 | 1.715 | 1.537 | 2.089 | 1.786 | 1.726 | 1.775 | 1.811 | 0.222 | 0.205 |
| Cytidine 5-triphosphate | 4.760 | 5.323 | 6.424 | 5.988 | 5.364 | 5.844 | 7.162 | 7.760 | 6.520 | 5.541 | 7.604 | 7.849 | 5.617 | 7.073 | 0.587 | 0.896 |
| Deoxycytidine 5-triphosphate | 0.080 | 0.073 | 0.085 | 0.089 | 0.083 | 0.082 | 0.151 | 0.148 | 0.124 | 0.125 | 0.134 | 0.131 | 0.082 | 0.136 | 0.005 | 0.011 |
| Deoxythymidine 5-triphosphate | 0.378 | 0.425 | 0.501 | 0.481 | 0.417 | 0.422 | 0.629 | 0.645 | 0.519 | 0.425 | 0.644 | 0.630 | 0.437 | 0.582 | 0.045 | 0.090 |
| Dihydroorotic acid | 0.188 | 0.161 | 0.190 | 0.257 | 0.198 | 0.149 | 0.107 | 0.102 | 0.114 | 0.096 | 0.123 | 0.103 | 0.190 | 0.107 | 0.038 | 0.010 |
| Dihydroxyacetone phosphate | 29.116 | 23.912 | 21.909 | 30.115 | 26.045 | 18.372 | 18.928 | 12.795 | 20.991 | 11.401 | 10.787 | 11.788 | 24.912 | 14.448 | 4.446 | 4.368 |
| Flavin adenine dinucleotide | 0.176 | 0.195 | 0.221 | 0.256 | 0.192 | 0.225 | 0.264 | 0.274 | 0.221 | 0.290 | 0.294 | 0.299 | 0.211 | 0.274 | 0.029 | 0.029 |
| Fructose 1,6-biphosphate | 8.354 | 14.658 | 16.870 | 13.810 | 9.305 | 9.076 | 5.229 | 4.863 | 6.073 | 4.682 | 5.012 | 5.212 | 12.012 | 5.179 | 3.554 | 0.485 |
| Fructose 6-phosphate | 0.534 | 1.159 | 1.825 | 1.101 | 0.635 | 0.933 | 0.763 | 0.825 | 0.535 | 0.972 | 0.703 | 0.693 | 1.031 | 0.749 | 0.462 | 0.146 |
| gamma-Glu-Cys | 0.363 | 0.423 | 0.456 | 0.448 | 0.379 | 0.421 | 0.477 | 0.388 | 0.302 | 0.447 | 0.481 | 0.442 | 0.415 | 0.423 | 0.037 | 0.068 |
| Gluconic acid | 0.300 | 0.309 | 0.307 | 0.336 | 0.329 | 0.303 | 0.157 | 0.147 | 0.129 | 0.146 | 0.146 | 0.161 | 0.314 | 0.148 | 0.015 | 0.011 |
| Glucose 6-phosphate | 2.867 | 4.605 | 7.233 | 4.344 | 2.782 | 4.445 | 4.710 | 3.649 | 3.547 | 4.411 | 3.645 | 3.921 | 4.379 | 3.980 | 1.615 | 0.476 |
| Glutamic acid | 99.706 | 77.960 | 121.516 | 113.306 | 102.859 | 106.170 | 107.889 | 94.560 | 111.916 | 82.946 | 80.825 | 85.485 | 103.586 | 93.937 | 14.798 | 13.285 |
| Glutamine | 289.607 | 236.141 | 391.940 | 351.360 | 302.566 | 351.336 | 302.209 | 285.194 | 345.646 | 240.736 | 237.921 | 252.967 | 320.492 | 277.446 | 55.496 | 42.032 |
| Glutathione (oxidized) | 1.240 | 0.888 | 0.836 | 1.178 | 1.070 | 0.770 | 0.737 | 0.700 | 0.788 | 0.732 | 0.788 | 0.645 | 0.997 | 0.732 | 0.193 | 0.055 |
| Glutathione (reduced) | 31.141 | 30.969 | 31.397 | 34.304 | 32.114 | 32.327 | 32.507 | 33.001 | 30.432 | 31.515 | 32.792 | 33.641 | 32.042 | 32.315 | 1.231 | 1.156 |
| Glyceric acid | 0.155 | 0.293 | 0.288 | 0.203 | 0.241 | 0.196 | 0.193 | 0.151 | 0.161 | 0.302 | 0.177 | 0.227 | 0.229 | 0.202 | 0.055 | 0.056 |
| Guanine | 0.281 | 0.404 | 0.427 | 0.428 | 0.374 | 0.382 | 0.325 | 0.293 | 0.376 | 0.491 | 0.292 | 0.350 | 0.383 | 0.354 | 0.054 | 0.075 |
| Guanosine | 0.373 | 0.674 | 0.651 | 0.617 | 0.579 | 0.404 | 0.281 | 0.247 | 0.352 | 0.356 | 0.232 | 0.307 | 0.550 | 0.296 | 0.129 | 0.052 |
| Guanosine 5-diphosphate | 4.429 | 1.156 | 4.838 | 4.720 | 4.276 | 4.598 | 7.102 | 5.799 | 4.560 | 6.115 | 6.158 | 5.064 | 4.003 | 5.800 | 1.409 | 0.894 |
| Guanosine 5-triphosphate | 14.293 | 14.817 | 17.986 | 16.720 | 15.403 | 12.352 | 20.907 | 16.295 | 15.122 | 6.952 | 18.693 | 18.131 | 15.262 | 16.017 | 1.958 | 4.870 |
| Hexoses | 251.062 | 436.511 | 434.636 | 272.171 | 471.846 | 153.754 | 199.312 | 124.687 | 245.348 | 216.370 | 157.264 | 233.885 | 336.663 | 196.145 | 128.674 | 46.653 |
| Histidine | 5.798 | 5.013 | 6.443 | 5.794 | 5.147 | 7.123 | 8.827 | 9.830 | 9.918 | 9.155 | 10.179 | 9.652 | 5.887 | 9.593 | 0.796 | 0.508 |
| Hydroxyglutaric acid | 0.206 | 0.200 | 0.195 | 0.232 | 0.227 | 0.187 | 0.204 | 0.198 | 0.176 | 0.199 | 0.209 | 0.208 | 0.208 | 0.199 | 0.018 | 0.012 |
| Hypotaurine | 0.501 | 0.305 | 0.404 | 0.543 | 0.353 | 0.496 | 0.334 | 0.377 | 0.335 | 0.323 | 0.531 | 0.302 | 0.434 | 0.367 | 0.094 | 0.084 |
| Inosine | 3.849 | 4.388 | 4.376 | 4.886 | 3.920 | 3.534 | 4.287 | 4.345 | 4.328 | 5.355 | 3.783 | 4.054 | 4.159 | 4.359 | 0.484 | 0.533 |
| Inosine 5-monophosphate | 0.223 | 0.266 | 0.284 | 0.235 | 0.196 | 0.203 | 0.161 | 0.151 | 0.131 | 0.200 | 0.237 | 0.212 | 0.234 | 0.182 | 0.035 | 0.041 |
| Isoleucine | 45.622 | 51.944 | 67.235 | 52.861 | 52.941 | 47.751 | 41.990 | 59.960 | 65.777 | 69.751 | 70.376 | 60.412 | 53.059 | 61.378 | 7.559 | 10.481 |
| Kynurenine | 0.183 | 0.247 | 0.293 | 0.245 | 0.231 | 0.233 | 0.346 | 0.341 | 0.304 | 0.391 | 0.360 | 0.387 | 0.239 | 0.355 | 0.035 | 0.032 |
| Lactic acid | 249.018 | 363.168 | 457.203 | 304.668 | 404.897 | 229.095 | 345.647 | 257.233 | 410.806 | 333.899 | 242.740 | 332.435 | 334.675 | 320.460 | 89.600 | 61.901 |
| Leucine | 41.010 | 54.441 | 63.483 | 50.970 | 44.979 | 48.134 | 70.073 | 63.250 | 61.163 | 68.146 | 58.930 | 62.257 | 50.503 | 63.970 | 7.880 | 4.276 |
| Malic acid | 5.645 | 5.775 | 6.314 | 6.632 | 5.707 | 4.372 | 3.011 | 2.523 | 2.924 | 2.531 | 2.153 | 2.346 | 5.741 | 2.581 | 0.776 | 0.331 |
| Methionine | 16.676 | 21.487 | 27.125 | 20.577 | 20.063 | 21.458 | 25.334 | 22.572 | 21.877 | 24.389 | 22.740 | 22.264 | 21.231 | 23.196 | 3.388 | 1.356 |
| N-Acetylaspartic acid | 0.006 | 0.009 | 0.008 | 0.007 | 0.007 | 0.007 | 0.008 | 0.009 | 0.007 | 0.008 | 0.009 | 0.008 | 0.007 | 0.008 | 0.001 | 0.001 |
| N-Acetylglutamic acid | 0.025 | 0.025 | 0.030 | 0.030 | 0.028 | 0.025 | 0.028 | 0.029 | 0.025 | 0.029 | 0.029 | 0.029 | 0.027 | 0.028 | 0.003 | 0.002 |
| N-Acetylneuraminic acid | 3.074 | 3.656 | 4.107 | 3.475 | 2.847 | 3.525 | 6.379 | 6.409 | 5.590 | 6.491 | 6.129 | 5.955 | 3.447 | 6.159 | 0.444 | 0.343 |
| O-Phosphorylethanolamine | 1.400 | 1.086 | 1.581 | 1.717 | 1.230 | 1.600 | 1.368 | 1.290 | 1.204 | 1.432 | 1.263 | 1.159 | 1.436 | 1.286 | 0.242 | 0.102 |
| Orotic acid | 0.167 | 0.137 | 0.152 | 0.202 | 0.198 | 0.143 | 0.149 | 0.119 | 0.117 | 0.099 | 0.102 | 0.127 | 0.167 | 0.119 | 0.028 | 0.018 |
| Pantothenic acid | 2.487 | 2.966 | 3.188 | 3.305 | 3.003 | 2.756 | 3.811 | 3.666 | 3.378 | 3.820 | 3.841 | 4.108 | 2.951 | 3.771 | 0.296 | 0.240 |
| Phenylalanine | 23.135 | 30.848 | 37.035 | 28.529 | 27.048 | 28.360 | 41.207 | 36.871 | 36.474 | 39.300 | 37.496 | 36.339 | 29.159 | 37.948 | 4.619 | 1.927 |
| Phenylpyruvic acid | 1.782 | 3.796 | 3.694 | 2.248 | 3.213 | 1.816 | 2.843 | 2.178 | 2.829 | 3.371 | 2.659 | 3.181 | 2.758 | 2.844 | 0.923 | 0.417 |
| Proline | 17.909 | 21.920 | 27.112 | 21.862 | 18.618 | 32.001 | 40.069 | 41.747 | 34.559 | 39.995 | 38.525 | 34.219 | 23.237 | 38.186 | 5.388 | 3.114 |
| Pyridoxal | 0.016 | 0.028 | 0.030 | 0.021 | 0.024 | 0.015 | 0.024 | 0.019 | 0.023 | 0.026 | 0.021 | 0.024 | 0.022 | 0.023 | 0.006 | 0.003 |
| Pyridoxine | 0.284 | 0.612 | 0.550 | 0.378 | 0.519 | 0.252 | 0.370 | 0.307 | 0.360 | 0.448 | 0.365 | 0.458 | 0.433 | 0.385 | 0.149 | 0.058 |
| Pyroglutamic acid | 28.656 | 50.427 | 48.356 | 32.294 | 42.959 | 23.678 | 29.590 | 23.352 | 31.397 | 36.114 | 26.610 | 32.778 | 37.728 | 29.974 | 11.053 | 4.536 |
| Pyruvic acid | 8.340 | 12.116 | 13.927 | 9.433 | 11.559 | 7.549 | 5.833 | 4.324 | 4.716 | 5.599 | 4.030 | 5.058 | 10.488 | 4.927 | 2.449 | 0.707 |
| Riboflavin | 0.111 | 0.184 | 0.188 | 0.162 | 0.189 | 0.107 | 0.151 | 0.122 | 0.147 | 0.158 | 0.145 | 0.163 | 0.157 | 0.148 | 0.039 | 0.014 |
| Ribose 5-phosphate | 0.418 | 0.504 | 0.563 | 0.454 | 0.412 | 0.331 | 0.352 | 0.236 | 0.336 | 0.307 | 0.195 | 0.257 | 0.447 | 0.281 | 0.081 | 0.061 |
| S-5-Adenosyl-L-homocysteine | 0.672 | 0.935 | 0.869 | 1.008 | 0.904 | 0.682 | 0.638 | 0.561 | 0.682 | 0.990 | 0.705 | 0.649 | 0.845 | 0.704 | 0.138 | 0.148 |
| Sedoheptulose-7-phosphate | 0.089 | 0.160 | 0.201 | 0.152 | 0.117 | 0.097 | 0.069 | 0.076 | 0.087 | 0.096 | 0.047 | 0.087 | 0.136 | 0.077 | 0.043 | 0.017 |
| Serine | 14.576 | 12.883 | 14.741 | 15.459 | 13.007 | 17.955 | 18.075 | 18.477 | 17.396 | 17.709 | 18.086 | 16.330 | 14.770 | 17.679 | 1.861 | 0.757 |
| Succinic acid | 1.553 | 2.109 | 2.070 | 1.807 | 2.072 | 1.393 | 1.171 | 0.975 | 1.000 | 1.553 | 1.256 | 1.254 | 1.834 | 1.202 | 0.304 | 0.211 |
| Taurine | 159.612 | 112.924 | 160.666 | 169.485 | 159.866 | 146.714 | 140.176 | 119.464 | 132.232 | 100.657 | 100.443 | 105.222 | 151.544 | 116.366 | 20.270 | 17.045 |
| Thiamine | 0.312 | 0.500 | 0.508 | 0.371 | 0.507 | 0.300 | 0.373 | 0.293 | 0.365 | 0.455 | 0.327 | 0.490 | 0.416 | 0.384 | 0.100 | 0.075 |
| Threonine | 169.866 | 141.301 | 225.780 | 201.053 | 173.129 | 216.022 | 254.504 | 229.749 | 257.396 | 191.417 | 187.099 | 195.660 | 187.859 | 219.304 | 31.980 | 32.165 |
| Thymidine 5-diphosphate | 0.405 | 0.317 | 0.371 | 0.438 | 0.356 | 0.409 | 0.543 | 0.534 | 0.483 | 0.563 | 0.542 | 0.503 | 0.382 | 0.528 | 0.044 | 0.029 |
| Thymidine 5-monophosphate | 0.085 | 0.097 | 0.096 | 0.112 | 0.086 | 0.076 | 0.054 | 0.051 | 0.053 | 0.057 | 0.056 | 0.060 | 0.092 | 0.055 | 0.013 | 0.003 |
| Tryptophan | 25.077 | 30.630 | 38.894 | 29.930 | 28.365 | 32.617 | 46.336 | 44.571 | 41.825 | 49.795 | 44.952 | 43.758 | 30.919 | 45.206 | 4.652 | 2.696 |
| Tyrosine | 32.131 | 28.867 | 46.213 | 37.432 | 36.393 | 39.603 | 58.514 | 50.855 | 60.678 | 40.384 | 40.743 | 43.270 | 36.773 | 49.074 | 6.029 | 9.006 |
| UDP-Glucose / UDP-Galactose | 10.993 | 13.101 | 18.004 | 13.864 | 12.305 | 14.957 | 14.795 | 13.624 | 12.940 | 13.651 | 14.035 | 12.961 | 13.871 | 13.668 | 2.432 | 0.698 |
| UDP-Glucuronic Acid | 3.353 | 3.894 | 4.310 | 4.322 | 3.876 | 4.046 | 3.760 | 3.637 | 3.007 | 3.439 | 3.910 | 3.606 | 3.967 | 3.560 | 0.358 | 0.313 |
| Uracil | 0.024 | 0.051 | 0.048 | 0.036 | 0.043 | 0.029 | 0.020 | 0.022 | 0.031 | 0.038 | 0.031 | 0.031 | 0.038 | 0.029 | 0.011 | 0.007 |
| Uric acid | 3.629 | 6.190 | 7.232 | 4.642 | 5.702 | 4.191 | 4.721 | 3.908 | 4.377 | 5.209 | 4.568 | 4.776 | 5.264 | 4.593 | 1.352 | 0.435 |
| Uridine | 0.671 | 0.932 | 0.955 | 0.956 | 0.841 | 0.688 | 0.576 | 0.545 | 0.591 | 0.719 | 0.542 | 0.598 | 0.840 | 0.595 | 0.132 | 0.065 |
| Uridine 5-diphosphate | 5.741 | 4.567 | 5.948 | 6.830 | 4.612 | 6.934 | 7.763 | 6.910 | 6.357 | 6.958 | 7.039 | 6.193 | 5.772 | 6.870 | 1.030 | 0.558 |
| Uridine 5-monophosphate | 5.104 | 4.791 | 5.501 | 5.678 | 4.839 | 4.508 | 4.416 | 3.912 | 3.866 | 4.725 | 3.876 | 3.568 | 5.070 | 4.061 | 0.448 | 0.425 |
| Uridine 5-triphosphate | 38.759 | 40.221 | 55.717 | 44.954 | 43.021 | 50.352 | 56.986 | 52.112 | 44.717 | 72.576 | 48.748 | 48.708 | 45.504 | 53.974 | 6.445 | 9.993 |
| Valine | 43.377 | 42.273 | 70.327 | 53.589 | 48.705 | 51.855 | 82.028 | 70.576 | 88.847 | 64.588 | 58.645 | 62.420 | 51.688 | 71.184 | 10.175 | 11.895 |
| Xanthine | 0.700 | 1.310 | 1.320 | 0.985 | 0.675 | 1.308 | 0.782 | 1.067 | 1.460 | 1.182 | 1.139 | 1.433 | 1.050 | 1.177 | 0.308 | 0.251 |
| Xanthosine | 0.027 | 0.044 | 0.045 | 0.038 | 0.035 | 0.033 | 0.021 | 0.017 | 0.025 | 0.028 | 0.019 | 0.033 | 0.037 | 0.024 | 0.007 | 0.006 |
| **Adenylate Energy Charge** | 0.322 | 0.390 | 0.397 | 0.354 | 0.368 | 0.329 | 0.360 | 0.359 | 0.338 | 0.269 | 0.325 | 0.353 | 0.360 | 0.334 | 0.031 | 0.035 |
| **GSH/GSSG** | 25.114 | 34.881 | 37.573 | 29.129 | 30.004 | 41.979 | 44.128 | 47.140 | 38.615 | 43.036 | 41.608 | 52.189 | 33.113 | 44.453 | 6.183 | 4.719 |

**Table S2: PD-L1 remodeling of the cancer cell metabolome.** Summary of unbiased metabolomics study performed on cell lysate and culture supernatant from TRAMP-C2 and TRAMP-C2-*Cd274*$^{-/-}$ after 48 hours in culture. All data in units of relative concentration, measured in μM. Culture supernatant data corrected using culture media without cells, to determine production/depletion of media metabolites.

**Table S3**. Summary of patient cohort for PD-L1 biomarker study.

| Sample ID | Sex | Age | Diagnosis | PD-L1 status | Average % infection |
|---|---|---|---|---|---|
| 203 | 62 | Female | lung squamous cell carcinoma | + | 0.5 |
| 204 | 83 | Female | endometrial cancer | - | 4.0 |
| 205 | 63 | Male | papillary renal cell carcinoma | + | 33.5 |
| 215 | 57 | Female | ovarian cancer | - | 0.9 |
| 221 | 67 | Female | endometrial cancer | + | 22.6 |
| 223 | 68 | Female | lung cancer | + | 27.9 |
| 224 | 70 | Female | hepatocellular carcinoma | - | 1.3 |
| 225 | 73 | Male | lung squamous cell carcinoma | + | 6.3 |
| 226 | 48 | Male | colon adenocarcinoma | - | 0.5 |
| 227 | 74 | Female | lung non-small cell carcinoma | - | 2.3 |
| 241 | 57 | Female | ovarian cancer | + | 0.2 |
| 242 | 73 | Female | chromophobe renal cell carcinoma | - | 0.1 |
| 247 | 62 | Female | colon adenocarcinoma | + | 9.3 |
| 248 | 63 | Female | ovarian cancer | - | 2.9 |
| 249 | 56 | Male | dermatofibrosarcoma protuberans | - | 0.2 |
| 253 | 62 | Male | neuroendocrine tumor | - | 3.8 |
| 257 | 67 | Female | uterine fibroid | + | 3.5 |
| 260 | 61 | Female | fibrous tumor of the abdomen | + | 11.4 |
| 283 | 52 | Female | breast carcinoma | + | 1.5 |
| 289 | 79 | Female | hepatic adenocarcinoma | - | 8.2 |
| 291 | 59 | Male | leiomyosarcoma | + | 0.1 |

**Table S3: Summary of patient cohort.**
